# Large-scale analysis of ligand binding mode similarities in the PDB using interaction fingerprints

**DOI:** 10.64898/2026.04.17.719144

**Authors:** Ibrahim Roshan Kunnakkattu, Preeti Choudhary, Adam Midlik, Jennifer R Fleming, Balakumaran Balasubramaniyan, Sreenath Sasidharan Nair, Sameer Velankar

## Abstract

Three-dimensional structures of protein-ligand complexes are essential for insights into the molecular principles that govern ligand recognition and binding. With more than 180,000 ligand-bound entries in the Protein Data Bank (PDB), representing over two million individual complexes, the volume of available structural data offers unprecedented opportunities for large-scale analysis of interaction patterns. Analysis of interaction patterns across the PDB archive can help discover similarities and differences in the binding modes of ligands, assisting in drug discovery. However, large-scale analysis of up-to-date information remains a significant challenge due to the rapid growth of data.

Here, we introduce the Extended Connectivity Interaction Fingerprint (ECIFP), an interaction-based fingerprint that simplifies 3D protein-ligand contact information into a fingerprint, while retaining key molecular and chemical features of the interacting fragments. The simpler fingerprint representation of the interaction data makes comparison of millions of protein-ligand complexes tractable. Benchmarking shows that ECIFP outperforms ligand-only Extended Connectivity Fingerprints in identifying similar binding sites across identical protein sequences occupied by chemically diverse ligands.

Our analysis showed that similarities calculated using ECIFP can be used to compare macromolecular complexes with similar or different ligands. In this study, we demonstrate two large-scale applications of ECIFP: (1) identification of distinct binding modes for over 9,000 ligands across the entire PDB, and (2) detection of binding-mode similarities among structurally diverse ligands within the same binding site across 48,870 binding sites from over 21,000 proteins.

## Introduction

The binding mode of a ligand in a protein-ligand complex refers to the specific spatial arrangement of the ligand within a protein’s binding site [1]. This includes the ligand’s position, orientation, and conformation, as well as its interactions with the protein [1]. Understanding the binding modes of ligands is crucial for deciphering the mechanism of molecular recognition and is routinely used to interpret structure-activity relationships.

Medicinal chemists visually inspect ligand binding modes to prioritise compounds for experimental validation in drug discovery projects [2]. Furthermore, they use existing knowledge of protein-ligand complexes to assist in determining the quality of ligands’ binding poses from docking studies [2]. The Protein Data Bank (PDB) is a key resource for mining the atomic details of ligand binding modes, currently holding over 240,000 experimentally determined macromolecular structures, of which over 75% are protein-ligand complexes [3]. Analysing the three dimensional structures of the protein-ligand complexes from the PDB provides insightful information like the shape of binding pockets, the composition of binding sites and key interactions between ligand and amino acids. Several studies have investigated the pattern of molecular interactions in the PDB and identified the frequency of common atomic interactions [4], spatial distribution of ligand atoms contacting amino acid residues [5] and similarities of ligand binding modes [6].

The multiple structures of the same protein available in the PDB present a great opportunity to investigate both conserved and specific structural properties of the protein and its interactions with ligands. PDBe-KB aggregated views of protein webpages facilitate the analysis of multiple structures of the same protein along with their structural, functional and biophysical annotations [7]. The superposed view of ligands available from these webpages helps to visualise different binding sites of a protein. However, for proteins with multiple binding sites and large numbers of ligands, it can be challenging to distinguish different binding sites by visual inspection. LIGYSIS webserver addresses this limitation to an extent by filtering out non-relevant ligands of a protein for its biological function and grouping ligands to different binding sites by clustering them based on their interaction patterns [8].

The calculated relative solvent accessibility, evolutionary divergence and enrichment in missense variation data of the binding sites helps to characterise the functional properties of the binding sites [8]. In addition to identifying functionally important binding sites, characterising the binding pattern is needed for understanding the specificity of a particular protein-ligand interaction. In case of binding sites with multiple ligands, comparing the binding patterns can uncover the complementarity of binding partners. sc-PDB database [9] provides features to search and compare binding modes of ligands based on their 3D interaction pattern, but this database was last updated in 2017, and the latest entries in the PDB are missing. The recently introduced PLINDER [10], a comprehensive collection of protein-ligand interaction systems, introduced several similarity measures to compare protein-ligand complexes. Although their utility was demonstrated for splitting training and test datasets for machine learning, their performance for binding mode comparisons was not investigated.

Existing methods in the literature for comparing ligand binding modes base their comparison on either the shape and orientation of ligands [11] or the interaction pattern between protein and ligands [6, 12]. Even though the methods based on the 3D shape and their structural alignments are interpretable, they are computationally expensive and therefore lack scalability. An alternative method commonly used for binding mode comparison is the representation of protein-ligand interaction patterns as interaction fingerprints (IFP). The simple representation of protein-ligand complexes as binary/integer fingerprints makes them amenable to large-scale comparisons. They have been widely used for comparison of docked ligand poses with native ligands in co-crystallised structures and have been incorporated into the scoring mechanism of different docking tools [13, 14].

Protein-ligand interaction fingerprints were pioneered by Structural Interaction Fingerprints (SIFt) proposed by Deng et al. [13]. SIFt represents protein-ligand interactions as a one-dimensional binary string based on seven predetermined interaction types [13]. Conversely, Structural Protein Ligand Interaction Fingerprints (SPLIF) uses an extended connectivity fingerprints (ECFP) atom hashing algorithm for encoding protein-ligand interactions instead of relying on predefined interaction types [14]. Similarly, Protein Ligand Extended Connectivity (PLEC) fingerprints encode protein–ligand interactions by pairing the ECFP environments from the ligand and the protein atoms [15]. Instead of encoding the 2D topology, several fingerprints use the pharmacophoric types of interacting atoms and fragments [16]. These wide varieties of interaction fingerprints (IFP) have also been applied for several use cases, including post-docking analysis [17, 18], virtual screening [19] and scaffold hopping [20]. Furthermore, some studies have explored interaction fingerprints (IFPs) for binding mode comparison of protein-ligand complexes. For example, Desaphy *et al.* applied Triplet Interaction Fingerprints (TIFPs) to align and compare protein–ligand complexes from the sc-PDB database [16]. In another approach, IFPs derived from 85 sequence-discontinuous residue positions in the KLIFS database were used to train machine-learning models for classifying different binding modes of kinase inhibitors [21]. Additionally, Kashara *et al.* introduced binding graphs to analyze protein–ligand complexes from the PDB, enabling the identification of similar ligand binding modes across structurally dissimilar proteins [6].

Although there have been several studies on the use of interaction based fingerprints, to the best of our knowledge, we have not found studies on their use for finding different binding modes of ligands from all the available 3D structures in the PDB. Specifically, we wanted to determine whether interaction fingerprints can discern the similarities and differences in the binding modes of ligands present in the same binding site of a protein collated from multiple PDB entries. To address these questions, we developed the Extended Connectivity Interaction Fingerprint (ECIFP), which encodes the interaction between proteins and ligands and uses them to compare binding modes of protein-ligand complexes. ECIFP creates a simple binary representation of protein-ligand complexes by encoding the interacting ligand and protein atoms and their environment. By restricting the encoding to only interacting fragments, ECIFP enables an easily interpretable pairwise comparison of protein−ligand complexes. To quantitatively assess its performance, we evaluated ECIFP for identifying similar binding sites using a benchmark dataset [22]. Furthermore, ECIFP was used to identify distinct ligand binding modes across the PDB and compare ligand binding modes within the same binding site of proteins.

## Results

### Overview of binding site and binding mode identification

ECIFP is an interaction-based fingerprint that encodes molecular features of interacting fragments of ligands and proteins. In this study, we used ECIFP to identify ligand binding modes by clustering protein–ligand complexes based on ECIFP fingerprint similarity scores, which range from 0 to 1, where 1 indicates maximum similarity. The interactions were calculated for all the protein-ligand complexes in the PDB using PDBe Arpeggio [23, 24].

Any non-polymer chemical component defined in one of the standard small molecule reference dictionaries - Chemical Component Dictionary (CCD) [25], Biologically Interesting molecule Reference Dictionary (BIRD) [26] or Covalently Linked Components (CLC) [27] were considered as ligands. Many small molecules in PDB structures are not biologically meaningful, because they are added as experimental necessities such as buffers, salts, cryoprotectants, or crystallisation agents [28, 29]. Hence, we removed such commonly known additives from our analysis [30, 31]. Additionally, we excluded protein-ligand complexes with low structural quality as assessed by the quality metrics reported in the wwPDB validation reports (see methods for details) [32, 33]. After generating ECIFP fingerprints for all selected protein-ligand complexes and clustering them based on fingerprint similarity, we systematically identified distinct binding modes of each ligand across the PDB. Furthermore, we clustered ligands bound to the same protein into distinct binding sites using their binding fingerprint similarity and compared their binding modes within each site.

### Binding modes of ligands across the PDB

Extended Connectivity Interaction Fingerprint (ECIFP) was used to capture the characteristic intermolecular interaction patterns of protein–ligand complexes in the PDB. ECIFP represents each protein-ligand complex using two interaction fingerprints - a ligand IFP and a protein IFP, both generated using Extended Connectivity Fingerprint (ECFP) atom-hashing algorithm (Figure 2). ECFP encodes the local atomic environment of each atom in a molecule based on the bond-radius neighbourhood. For example ECFP1 encodes the root atom and its direct neighbours. In this study, we used ECFP4, which captures atomic environments within a maximum radius of two bond steps. Importantly, only interacting atoms were considered as root atoms rather than all atoms in the molecule. As a result, the ligand IFP represents interacting ligand fragments, while the protein IFP represents interacting fragments of amino acid residues. This separation allows flexible weighting of ligand and protein contributions during pairwise similarity comparisons, depending on whether the analysis focuses on the same ligand across different binding sites or on different ligands within the same binding site. By restricting fingerprint generation to interacting atoms, ECIFP provides a compact yet informative representation of protein–ligand complexes that preserves essential molecular features.

**Figure 1.**
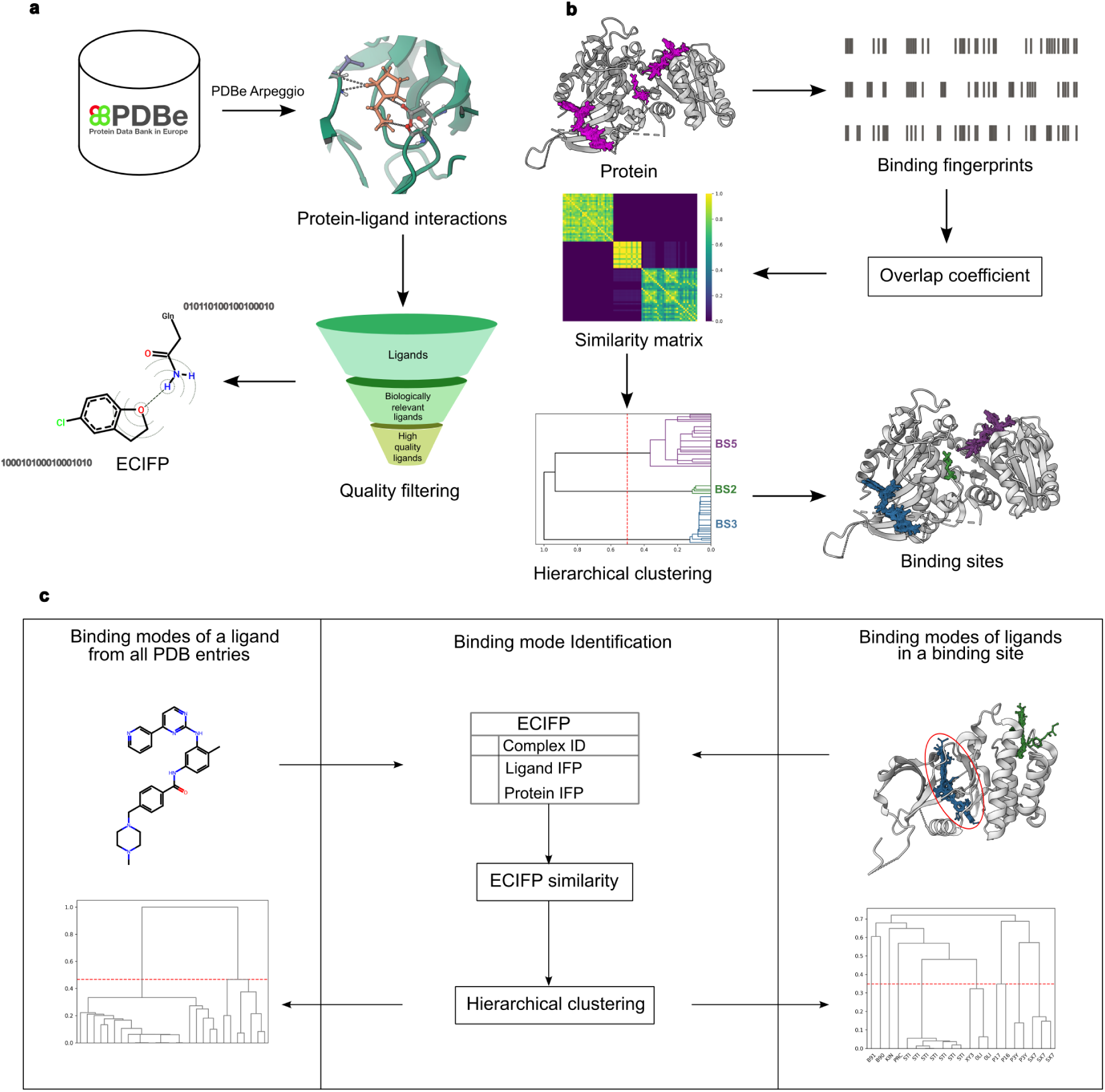
Key steps in binding site and binding mode identification. **a,** ECIFP calculation - protein-ligand interactions are calculated for all the PDB entries using PDBe Arpeggio, then filtered to include only high-quality biologically relevant ligands. ECIFP is generated for the filtered set of protein-ligand complexes and stored in a database table. **b,** Binding site identification - For all the ligands bound to the same protein, binding fingerprints are generated as the set of residue numbers of their interacting protein. The overlap coefficient was used to find similarities of binding fingerprints. Distinct binding sites are identified by hierarchical clustering of ligands, based on their binding fingerprint similarity. Ligands are coloured pink before clustering, after clustering all the ligands in the same cluster are coloured the same **c,** Binding mode identification - protein-ligand complexes are clustered based on the similarity of their ECIFPs to find their binding modes. The input protein-ligand complexes can correspond to either all the instances of a ligand from multiple PDB entries or multiple ligands from the same binding site of a protein. Ligands binding to the same site, for which the binding mode has been identified, are highlighted with a red ellipse.

**Figure 2.**
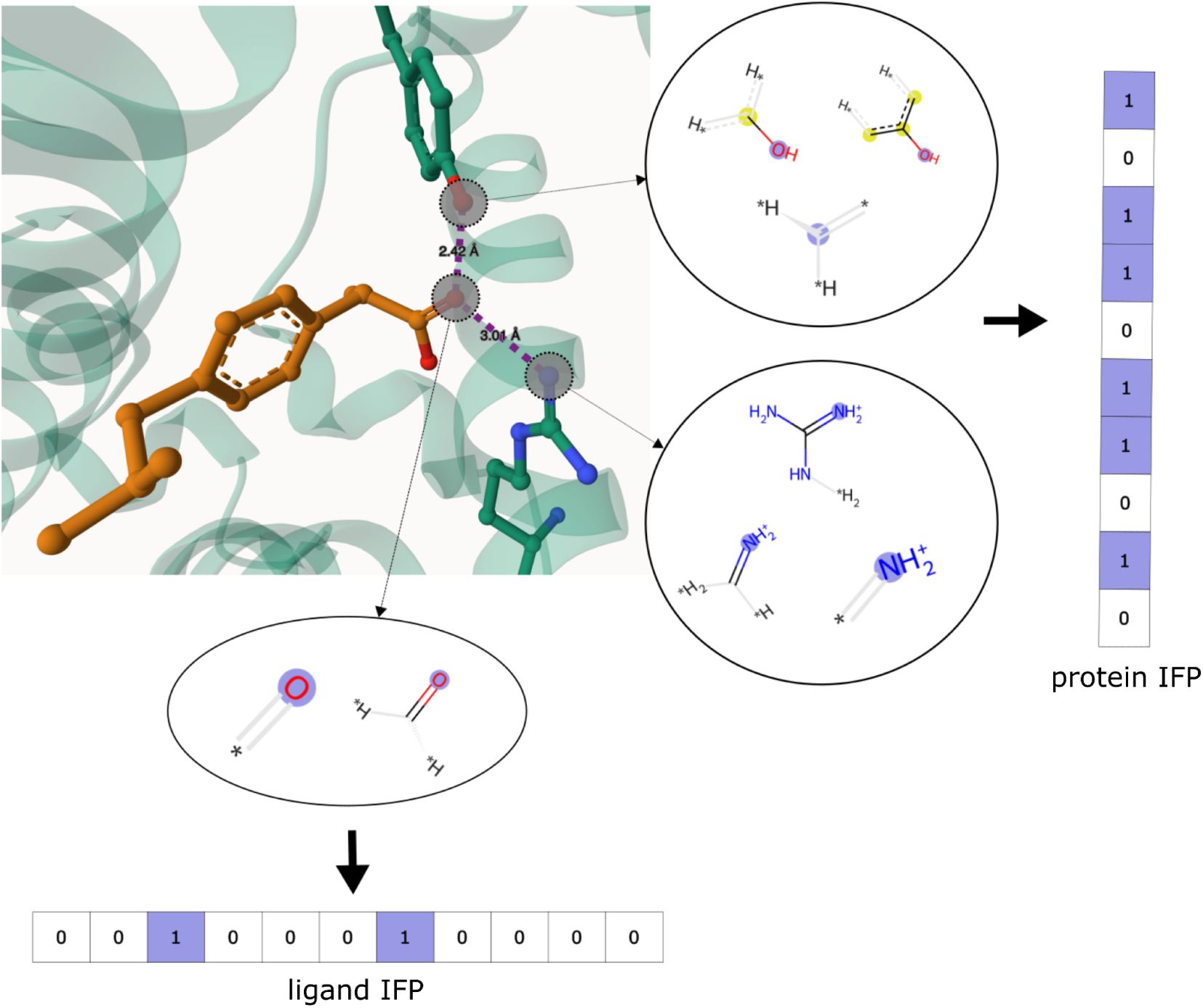
Interacting ligand and protein atoms are highlighted by a circle in the 3D view. The atomic environment of interacting atoms defined by ECFP4 algorithm is visualised using RDKit. The root atom is coloured in blue and aromatic atoms in the environment are colored in yellow. A binary interaction fingerprint is generated for both ligand and protein based on the atomic environments of their interacting atoms.

Clustering protein–ligand complexes based on similarity of these fingerprints revealed the distinct binding modes of ligands across the PDB. To restrict the analysis to biologically relevant ligands, salts, crystallisation agents, cryoprotectants, and ions were excluded (see Methods). After removing low quality protein-ligand models based on the ligand quality metrics in the wwwPDB validation report (see Quality filtering of protein-ligand complexes in Methods), binding modes were identified for 9,563 ligands with at least two complexes (instance of a ligand bound to a PDB entry). For the majority of ligands (75%), fewer than four complexes were available, and these ligands typically exhibited a single binding mode.

In contrast, several ligands were represented by a large number of complexes, with FAD, NAD, CLA, ADP, and HEC each observed in more than 4,000 complexes. The top ten ligands ranked by the number of binding modes are shown in Figure 3.

**Figure 3.**
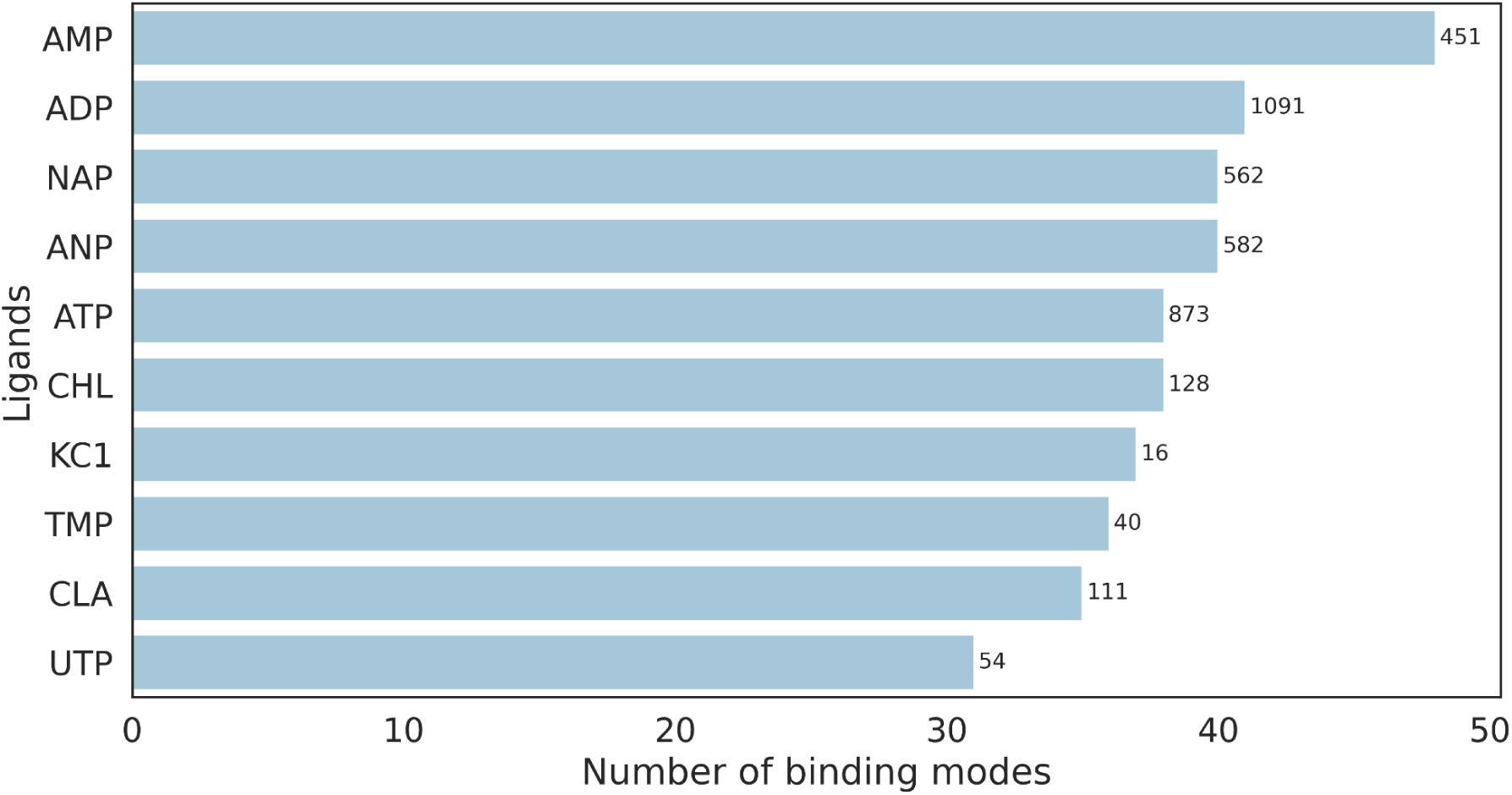
Top ten ligands based on number of binding modes.The number of binding modes of top ten ligands. Corresponding numbers of unique proteins (Uniprot IDs) are shown as bar labels.

Using minimax linkage method [34], which selects a prototype that minimises the maximum distance to any other data point in the cluster, we were able to identify representative binding modes for each cluster. This is particularly useful for generating a representative data set of ligand binding modes when a ligand has many protein-ligand complexes, enabling efficient visual inspection. For instance, three distinct clusters with their own representative binding modes were identified for the broad-based retroviral protease inhibitor TL-3 [35] (CCD ID: 3TL) from 30 protein-ligand models (Figure 4). Cluster one contained feline immunodeficiency virus proteases (FIV PR) with the carboxybenzyl groups of TL-3 at P4 and P4’ located along the major axis of the active site [36]. In contrast, cluster two contained human immunodeficiency virus proteases (HIV PR), where these groups were oriented over the flap region [36]. Notably, this cluster also included one FIV PR structure (PDB ID: 2hah) in which 12 active-site residues were substituted with their HIV PR equivalents, consistent with its reported similarity in interaction pattern to HIV PR [35]. Finally, cluster three consisted of a single structure (PDB ID: 3slz) corresponding to the xenotropic murine leukemia virus-related virus protease which has a similar conformation of TL-3 as in FIV PR, however has more similarity to the interaction pattern of TL-3 in HIV PR (Supplementary Figure 2d).

**Figure 4.**
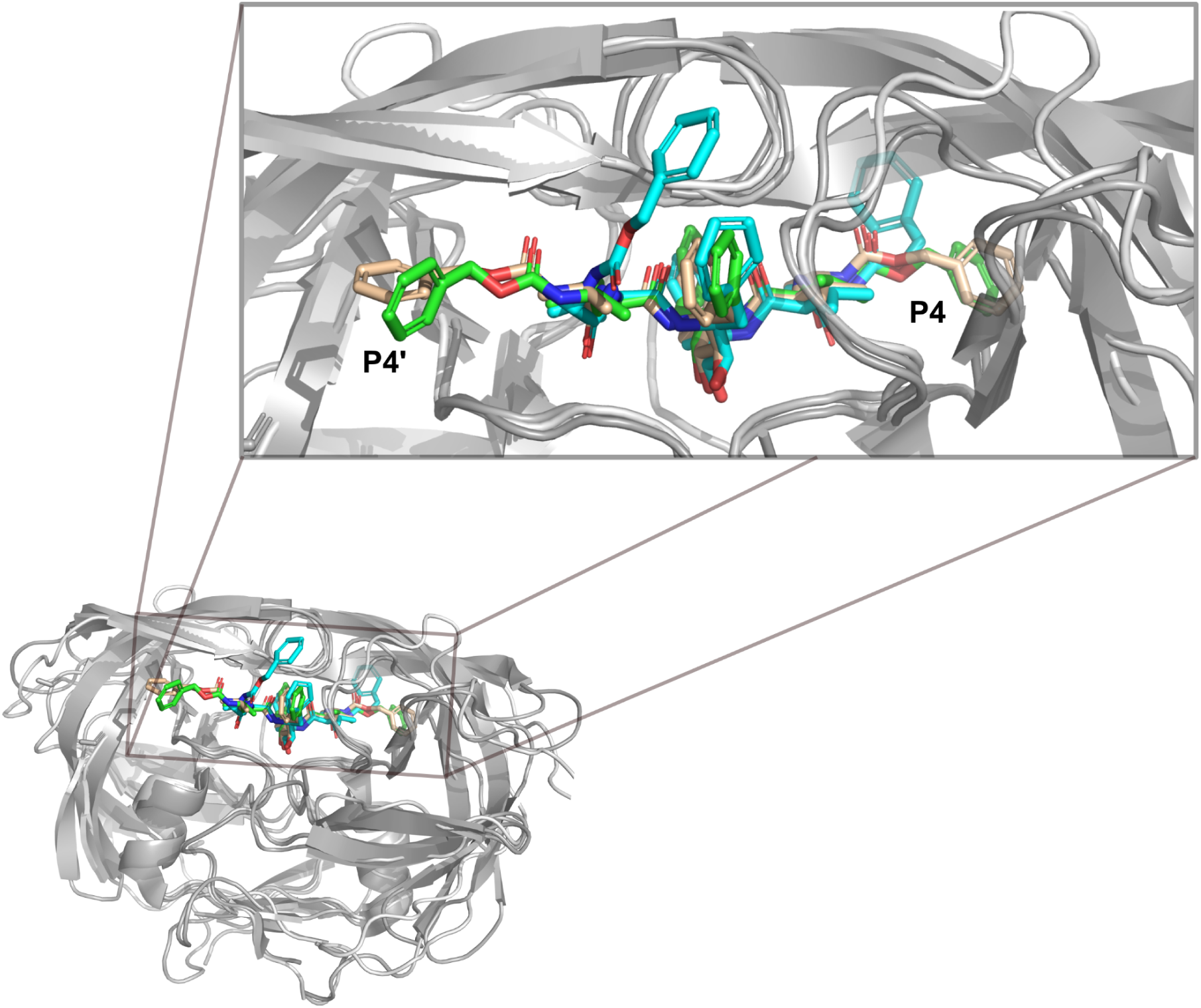
Representative binding modes of TL-3. Representative binding modes of TL-3 for each of the three clusters identified are shown after superposition of polymer chains. Binding modes of TL3 corresponding to cluster one, cluster two and cluster three are shown in green (1b11), cyan (4tvh) and tints of wheat colored (3slz) sticks.

### Cofactor binding modes are conserved among enzymes with similar function

Cofactors are ligands present in the active or allosteric sites of proteins and that are essential for their biological function. Many PDB entries contain cofactor-bound structures, and their analysis is crucial for a better understanding of protein function, such as enzyme mechanisms [37]. Here, we analysed the binding modes of 20 organic cofactors defined in the CoFactor database [38], by comparing the similarity of their ECIFPs. The number of complexes analysed for each cofactor and the corresponding number of binding modes identified are listed in Supplementary Table 1. To examine the relationship between cofactor binding modes and enzyme function, we binned the protein–cofactor complexes for each cofactor into uniformly spaced intervals of ECIFP similarity and, within each bin, computed the proportion of pairs that share the same EC number (Figure 5). For most cofactors, protein–cofactor pairs in high-similarity bins showed a high likelihood of sharing the same EC number, indicating that similar binding modes tend to correspond to similar enzymatic functions. An exception was HEME-A (HEA), which exhibited a high percentage of shared EC numbers across all similarity bins. This pattern was largely driven by the diverse binding modes of HEA observed in cytochrome c oxidase (Supplementary Figure 1).

**Figure 5.**
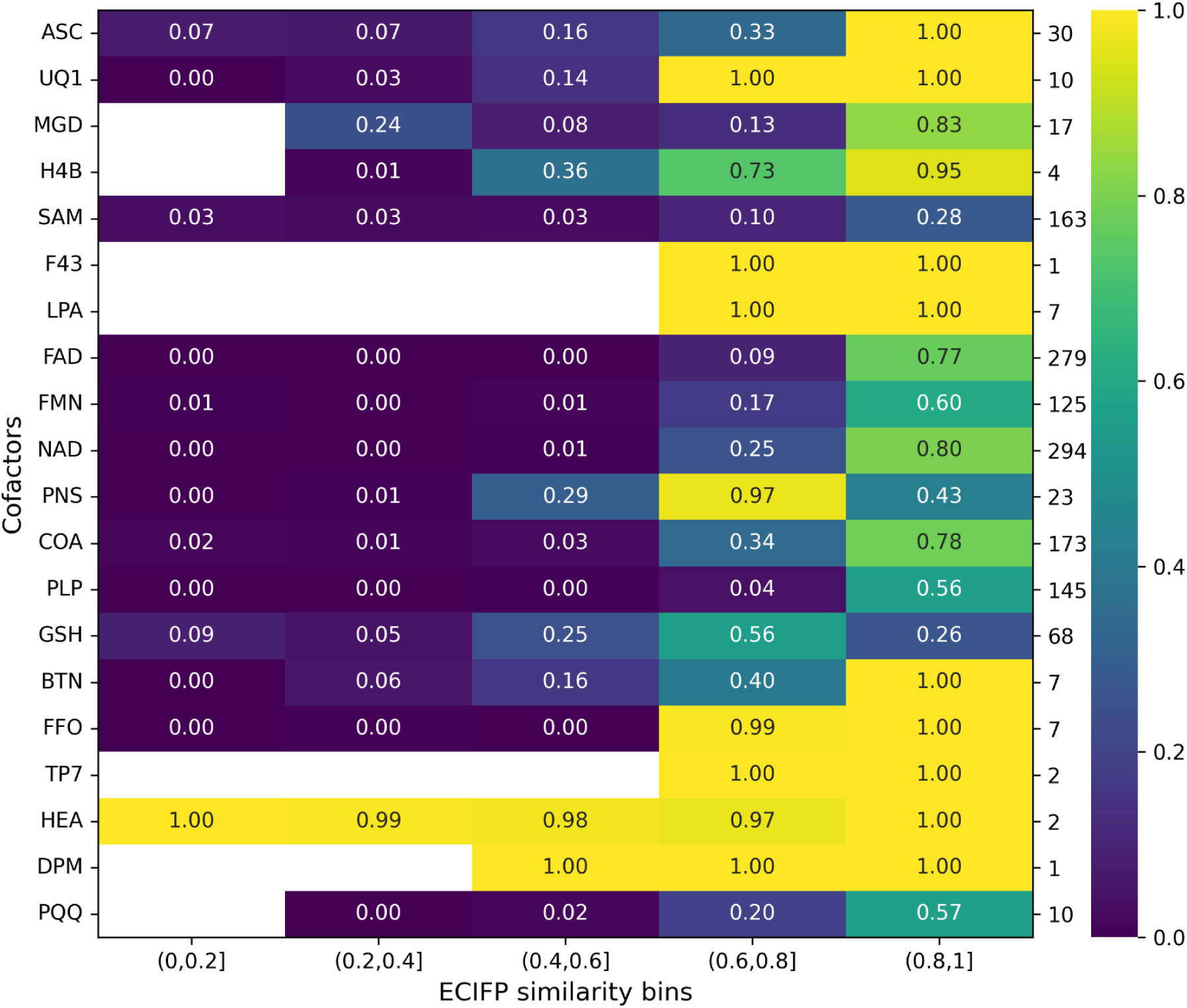
Fraction of protein-cofactor complexes sharing the same EC number in different similarity bins. Fraction of protein–cofactor complexes sharing the same EC number across different ECIFP similarity bins. Cofactors are labelled using their CCD IDs. For each cofactor, the number of unique EC numbers observed among all protein–cofactor pairs is shown on the secondary y-axis.

An illustrative example of using ECIFP to compare binding modes of cofactors is given in Figure 6. Here, we identified the two binding modes of S-adenosylmethionine (SAM) in the PDB entry 2c2b. The PDB entry 2c2b corresponds to the structure of threonine synthase, which is activated by the binding of SAM and catalyzes the ultimate step of threonine synthesis in *Arabidopsis thaliana* [39]. The crystal asymmetric unit has three dimers with two binding SAM molecules in the two activator binding sites located in the dimeric interface. The four bound SAM molecules give rise to two distinct binding modes (binding mode 1 - SAM 1 and SAM 2, binding mode 2 - SAM 3 and SAM 4) [39]. The superposition of the two chains using US-align [40] shows the perfect alignment of SAM1 and SAM3 as well as SAM2 and SAM4. ECIFP similarity scores were higher for SAM molecules with the same binding mode and lower for molecules in different binding modes (Figure 6 c)

**Figure 6.**
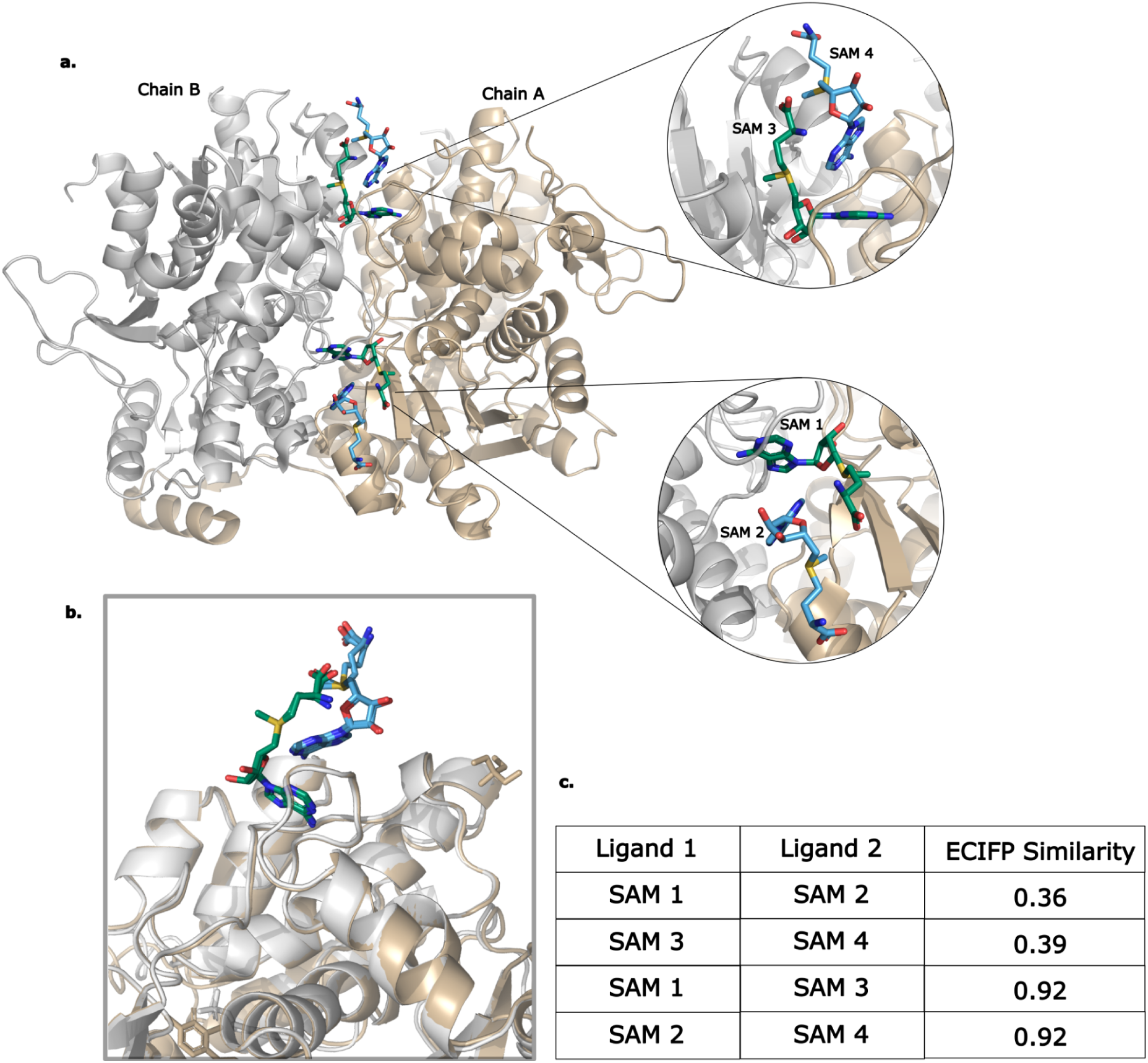
Binding modes of SAM in threonine synthase. **a**, Cartoon representation of threonine synthase dimer from the PDB entry 2C2B. Monomers represented by chain A and chain B are shown in tan and grey colours, respectively. The four SAM molecules at the dimer interface are in green (SAM 1 and SAM 3) or blue (SAM 2 and SAM 4) sticks. **b,** Superposition of chain A to chain B shows that corresponding SAM molecules in each binding site have similar binding modes. **c,** ECIFP similarities of SAM molecules at the dimer interface.

**Figure 7.**
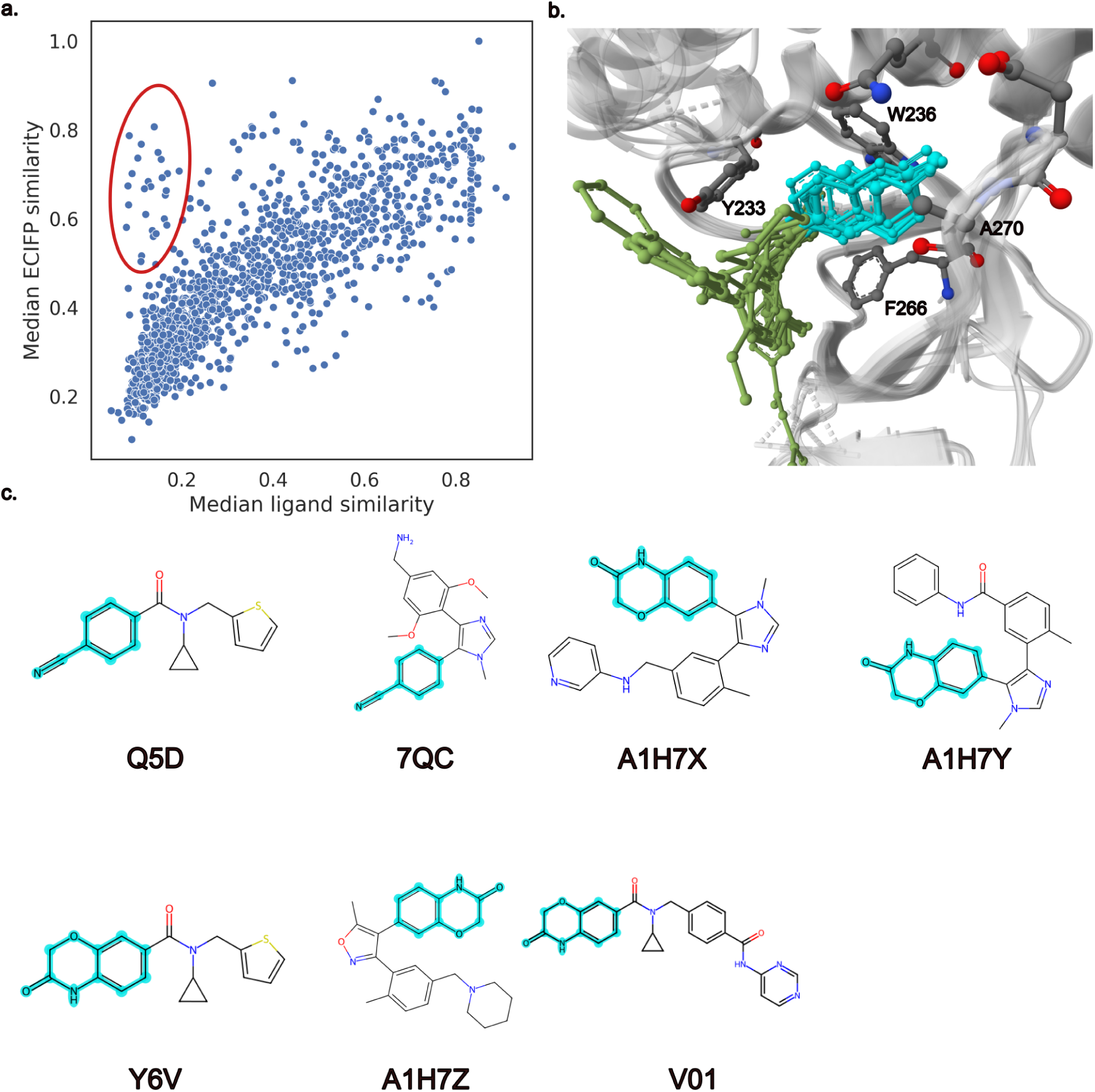
Binding modes of ligands in the same binding site. **a,** Scatter plot of median pairwise ligand similarities and median pairwise ECIFP similarities of ligands in the same binding site. Binding sites with low pairwise ligand similarities and high pairwise ECIFP similarities are highlighted in red ellipse. **b**, Binding modes of potent inhibitors of histone-lysine N-methyltransferase. All the ligands are oriented in a way the hydrogen bond acceptors (cyanophenyl group and benzoxazinone ring, coloured in cyan) form hydrogen bond to the backbone of Ala270 **c**, Potent inhibitors of histone-lysine N-methyltransferase labeled with their CCD ID. Cyanophenyl group and benzoxazinone ring are highlighted in cyan

**Figure 8.**
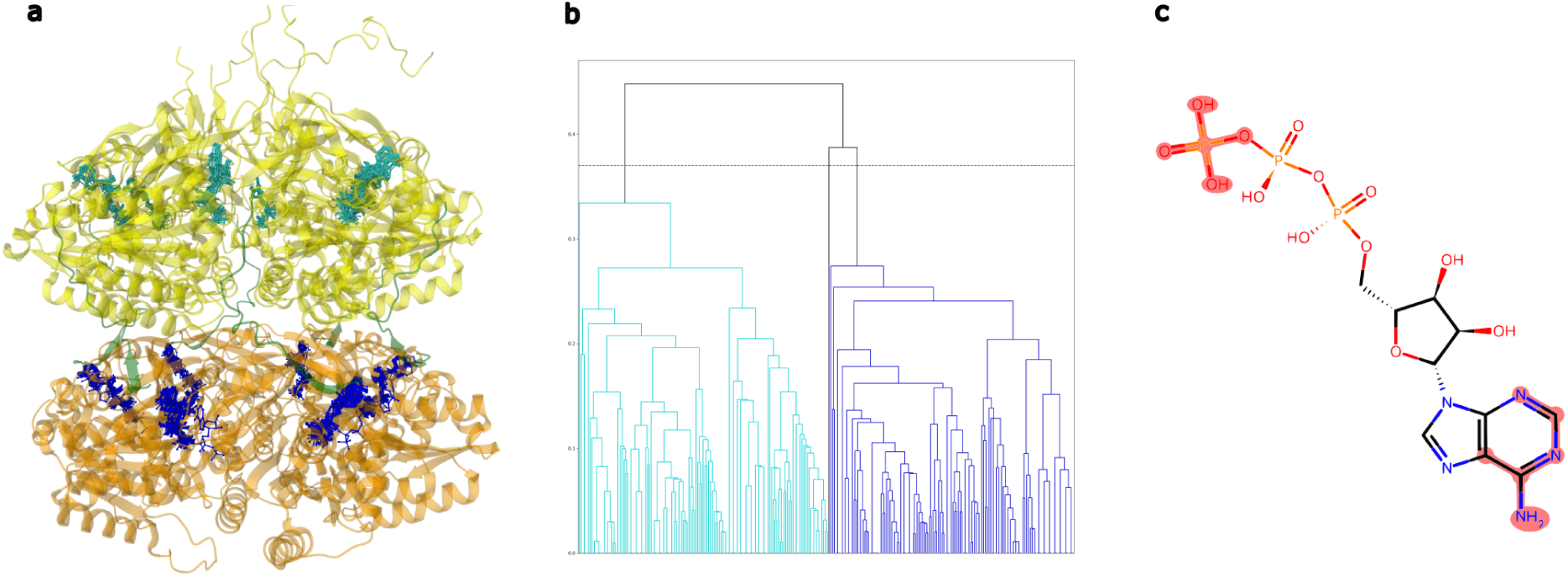
Binding sites and binding modes of ATP in circadian clock oscillator protein KaiC. **a,** Structure of KaiC homo-hexamer (PDB ID: 3dvl) after superposition of all the PDB entries mapping to the protein. N-terminal CI and C-terminal CII domains from individual subunits are coloured orange and yellow. The linkers between the domains are coloured green.

**Figure 9.**
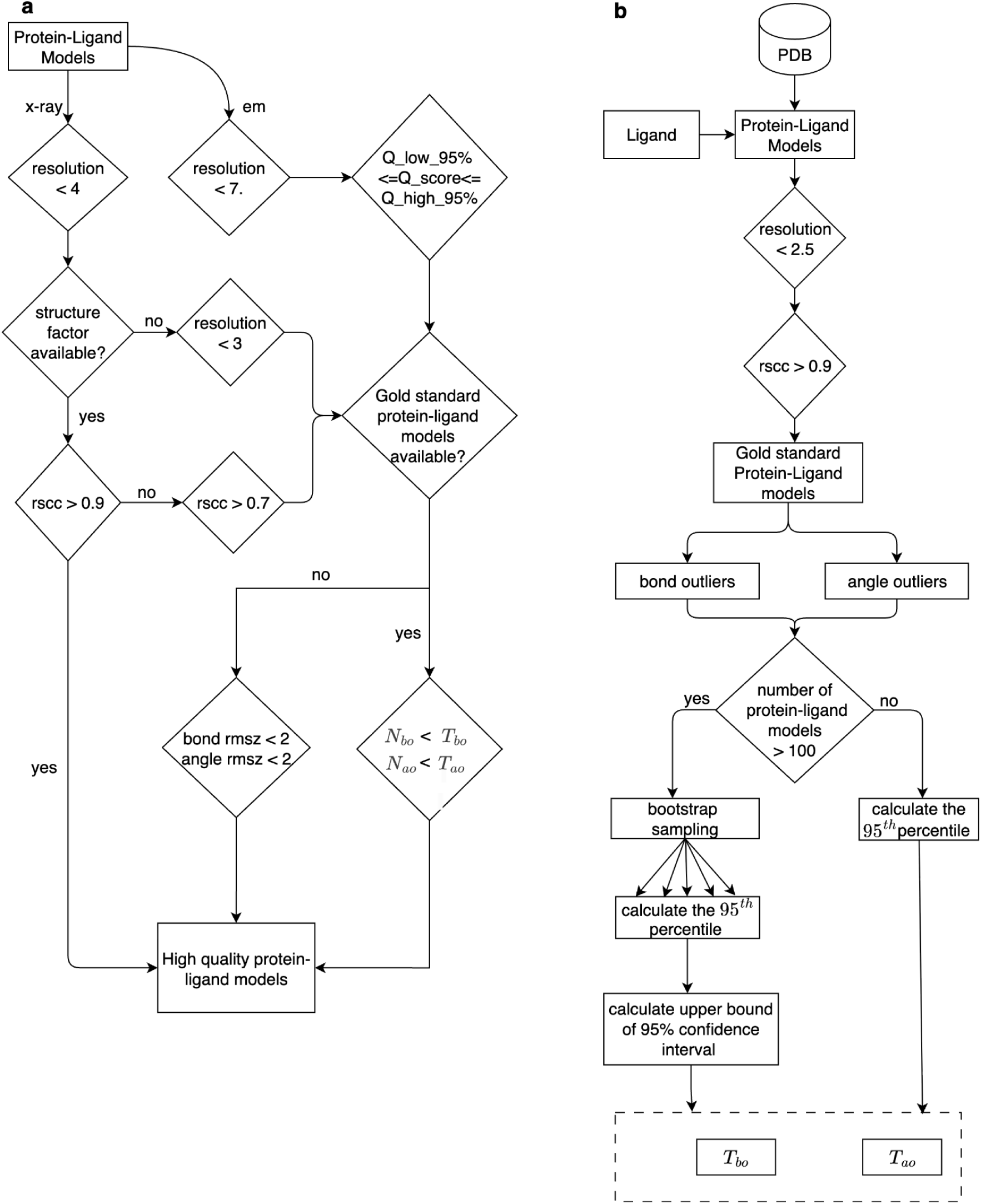
Pipeline for filtering protein-ligand models based on atomic model quality. **a,** Pipeline for filtering protein-ligand models based on atomic model quality metrics. N_bo_ - number of bond length outliers, N_bo_ - number of bond angle outliers, T_bo_ - maximum accepted number of bond length outliers, T_bo_ - maximum accepted number of bond angle outliers **B**, Pipeline for selecting a gold-standard set of protein-ligand models and defining cutoff values for the maximum number of accepted bond length and bond angle outliers

### ECIFP outperforms ligand-based fingerprints for binding-site similarity assessment

To quantitatively assess the protein-ligand complex similarities calculated by ECIFP, we evaluated its performance in identifying similar binding sites using a diverse benchmark data set - ProSPECCTs [22]. ProSPECCTs includes 10 objectively defined data sets of protein-ligand binding site pairs with classes defined as similar or dissimilar according to specific criteria. We limited our analysis to a subset of benchmark data sets, including “Structures with identical sequences” (DS1), “Kahraman data set” (DS2), “Barelier data set” (DS3) and “Data set of successful applications” (DS4) based solely on the availability of common protein-ligand complexes in ProSPECCTs and our data set. Each of these datasets were designed to evaluate multiple aspects of binding site comparison. DS1 contained structures with identical sequences, but bound to chemically different ligands [22]. The performance of a binding site similarity method on this dataset is a good measure of its ability to identify common properties of binding sites with a wide variety of ligands. DS2 and DS3 were derived from two previous studies [41, 42], and contain binding sites occupied by identical or highly similar ligands. DS2 is well-suited to test if methods can differentiate binding sites occupied by identical ligands from those occupied by different ligands. DS3 contained unrelated proteins bound to similar ligands, and hence, a good test set of methods for identifying local similarities of binding sites. Finally, DS4 was compiled by the authors of ProSPECCTs from known binding site similarities reported in the literature, and it is suited to test if methods are able to recover known binding site similarities within a set of diverse proteins [22]. The performance of ECIFP-based similarity metric was measured across these four data sets using area under receiver operating characteristic curve (AUC) (Table 2). It was also compared to a baseline model which used ligand similarity measured using the Tanimoto coefficients of 1024-bit ECFP4 fingerprints. Additionally, to visually inspect and evaluate the predictions, we identified a similarity cutoff of 0.3 based on the distribution of randomly selected protein-ligand model pairs (Supplementary Figure 3).

**Table 2.**
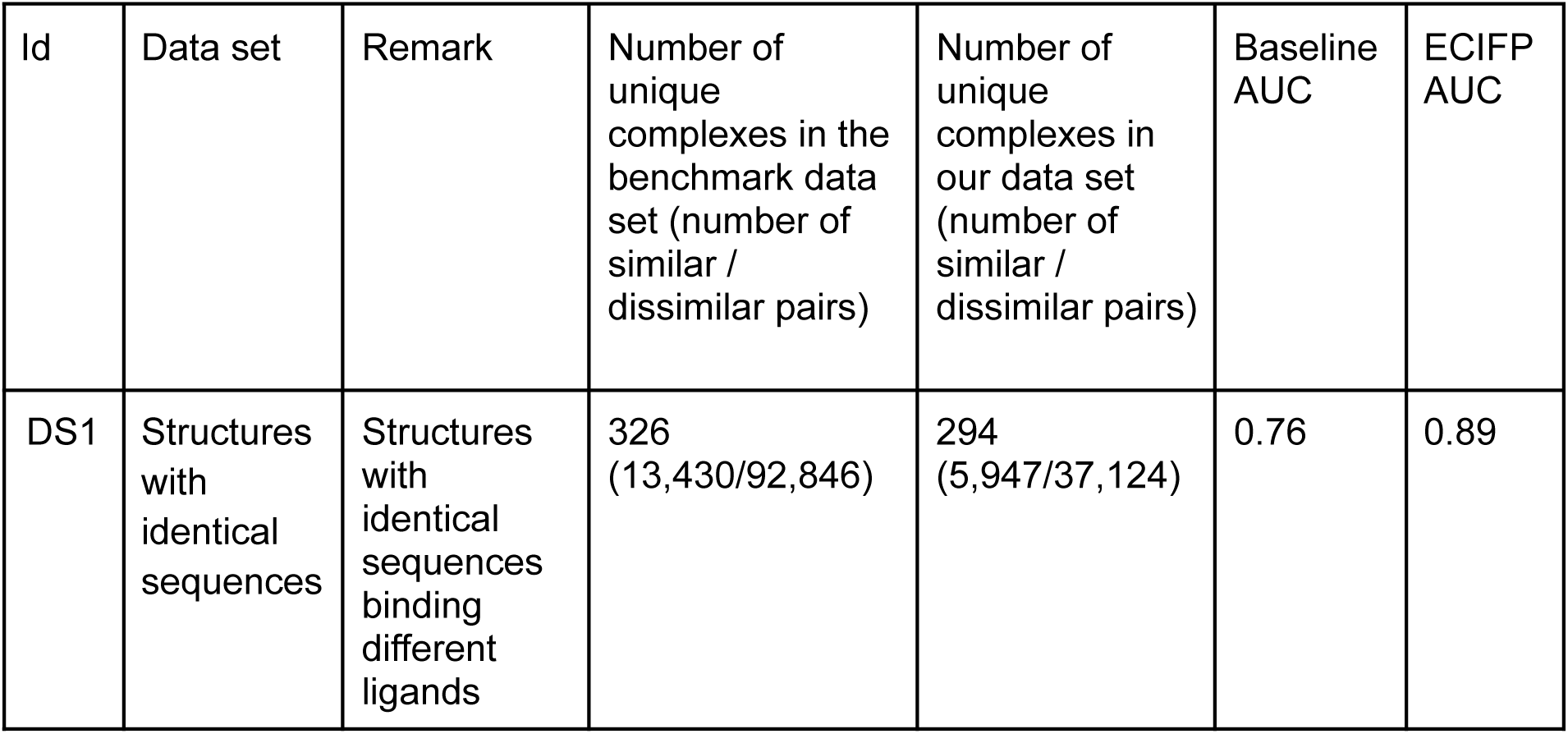
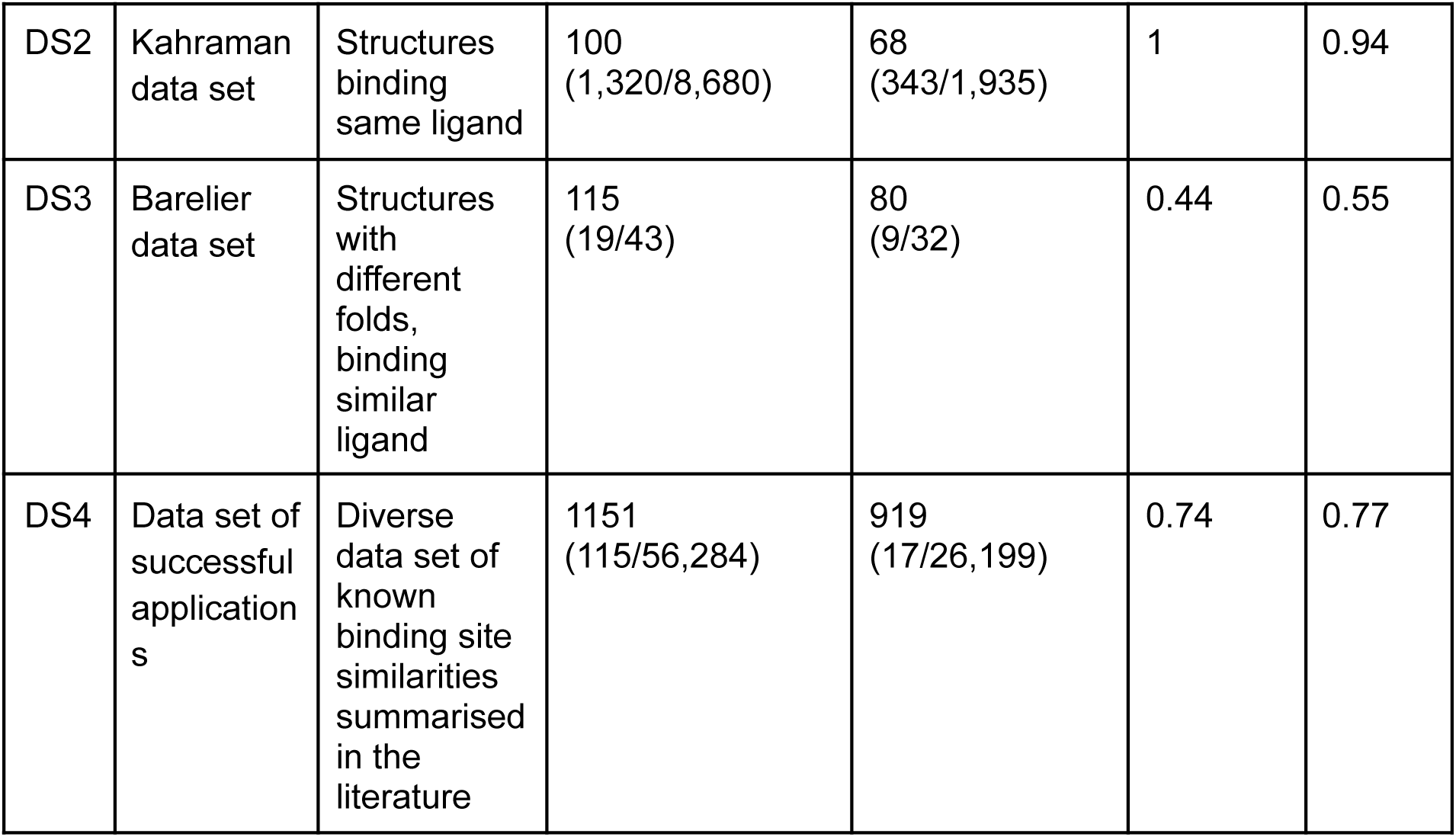
Evaluation of ECIFP similarities for binding site comparison.

Based on the AUC values, ECIFP similarities outperformed the baseline method in DS1. We found several pairs of similar binding sites with high ECIFP similarity even when the overall structural similarity of the ligand is low (Supplementary Table 3). In contrast, the baseline method had a perfect AUC of one in DS2. The high AUC scores can be primarily attributed to the design of this dataset rather than the effectiveness of the method to distinguish between binding sites. Similar binding sites were defined as those with identical ligands, and dissimilar binding sites as those with different ligands. However, similar, but not identical ligands can interact with the protein in similar ways, and we found several pairs like these in this data set (Supplementary Table 4). In contrast, we found 38 pairs of binding sites occupied by the same ligand, but had low ECIFP similarity. 84% of these pairs had phosphate as the ligand, and the low ECIFP similarity was mainly due to the difference in amino acids interacting with the phosphate. In DS3, the performance of ECIFP similarities was comparable to that of a random classifier, as was that of all the tools studied in ProSPECCTs [22]. DS3 was compiled by Barelier *et al.* [42] for investigating recognition of identical ligands by unrelated proteins. They grouped 59 ligands from 116 complexes into three groups - class A, class B and class C. Class A included pairs of complexes where the same ligand group made similar interactions with related binding site residues. In class B, the same ligand group interacted with different residues and in class C different ligand groups interacted with the protein. In ProSPECCTs data set, pairs from class A were classified as similar and others as different. Using an ECIFP similarity cutoff of 0.3, eight out of nine pairs from similar binding sites were correctly classified. However, sixteen out of twenty pairs of dissimilar binding sites were misclassified. Because the complexes in this dataset had the same ligand, they generally shared high similarity in their ligand IFP, primarily due to hydrophobic and aromatic interactions. The differences in the binding sites of the DS3 data set were defined by Barelier *et al.* based on visual inspection of key interactions and the physicochemical properties of the binding site local environment. They included water mediated interactions as well as interactions with other ligands in the binding site. Our analysis included only interactions between the ligand of interest and the amino acid residues. Moreover, all interaction types were weighted equally, and the similarity of ligand and protein IFPs were considered independently. These differences may have contributed to poor performance in classifying this data set. Finally, the performance of ECIFP similarities in DS4 was also moderate with an AUC comparable to the baseline method. Out of the 20 pairs of similar binding sites, eight of them had similar ligands, and they were correctly classified. However, the 12 pairs containing dissimilar ligands were misclassified. Several pairs of binding sites with class *dissimilar* in this data set had high ECIFP similarities. We manually analysed the superposition of 45 pairs of binding sites with ECIFP similarity greater than 0.6, and found that 34 of them had very good alignment in their binding sites (Supplementary Table 5). Additionally, there were 187 pairs of binding sites with ECIFP > 0.5. Most of them were ligands containing the nucleoside - adenosine. Adenosine has an adenine moiety and ribose sugar, both capable of forming hydrogen bonds. The N1 and N6 atoms of adenine can form hydrogen bonds to the backbone of amino acids and contribute to their recognition by different protein families and folds [43, 44]. Adenine can also form cation-pi and pi-pi interactions with aromatic amino acids. We found these interaction patterns to be shared among many protein-ligand modes in this dataset with high ECIFP similarity.

Overall, the proposed ECIFP-based similarity metric is able to identify similarities in the binding sites when there are some structural similarities in ligands occupying the binding sites. By encoding the interaction information, the method captures the similarities in interaction patterns even when the overall structural similarity between ligands is low.

### ECIFP identifies similar binding modes of ligands within the same binding site

As of February 2026, there are about 72,500 proteins (unique Uniprot IDs) whose sequence maps to one or more polymer sequences of PDB entries [45, 46]. For many proteins, there is more than one corresponding PDB entry with ligands - presenting an opportunity to investigate both conserved and specific structural features of proteins. Aggregating protein-ligand interactions from multiple PDB entries, we identified the distinct binding sites present in the proteins using the same approach as Utgés et al. [47]. Briefly, the method clusters ligands that interact with the same protein, based on the similarity of their binding fingerprints - defined as the set of UniProt residue numbers corresponding to ligand-interacting residues [8, 47, 48]. This avoided reliance on computationally expensive structural superpositions, while capturing spatially consistent sets of binding residues across multiple PDB entries of the same protein.

Using this approach, we identified 48,870 binding sites across 21,142 proteins (unique Uniprot IDs) corresponding to 102,843 PDB entries and 37,440 ligands, covering 54% of ligand bound structures in PDB. For 75% of proteins, fewer than two binding sites were identified. Furthermore, for 85% of all the pairs of binding sites, the minimum interatomic distance between ligands occupying different sites within the same protein exceeded 4 Å, indicating that binding sites identified using interaction fingerprints are well separated in three-dimensional space. The quality of binding site clustering was further assessed by visual inspection of the superposed structures of 20 randomly selected proteins (Supplementary Table 6). Except in one case, the identified binding sites were distinct and well separated in 3D space. In this exception (Beta-galactoside alpha-2,6-sialyltransferase 1, UniProt accession: P15907), two closely positioned sites were identified as distinct, as ligands at each site exhibit different binding modes and interact with different residues. Notably, alternate conformations of a single ligand (Cytidine-5′-monophosphate-5-N-acetylneuraminic acid; CCD ID: NCC) occupy both sites in two distinct binding modes.

To determine whether similar ligands bind to the proteins in similar modes, we compared the similarities of ligands in the same binding site to their binding mode similarity measured using ECIFP. Ligands were represented by ECFP4 fingerprints, and Tanimoto coefficients were used for similarity calculation. We only included the binding sites containing protein-ligand complexes with ECIFP, after quality filtering as described in the Methods. This resulted in 1,415 proteins and 1,565 binding sites. We compared the median pairwise ligand similarities with the median pairwise binding-mode similarities across binding sites, and observed a strong correlation between ligand similarity and ECIFP similarity (Pearson correlation coefficient = 0.81), indicating that structurally similar ligands tend to bind in similar modes at the same binding sites.

To evaluate whether ECIFP captures binding mode similarity beyond what is reflected by global ligand similarity, we calculated the coefficient of determination (R²), treating ECIFP similarity as the dependent variable and ligand similarity as the independent variable. The low R² value (0.04) indicates that most of the variance in ECIFP similarity cannot be explained by overall ligand similarity alone. This suggests that ECIFP encodes interaction-level features that are not directly tied to the full molecular structure. As illustrated in the scatterplot (Figure 4), several binding sites exhibit low median pairwise ligand similarity but high median pairwise ECIFP similarity. This highlights cases where structurally diverse ligands adopt similar binding modes by preserving key interaction motifs and spatial arrangements of functional fragments. One such example is shown in Figure 4 b, where chemically distinct potent inhibitors of histone-lysine N-methyltransferase (NSD2) bind in similar modes. Although their median and mean ligand similarities are low (0.26 and 0.28, respectively; SD = 0.16), the corresponding ECIFP similarities are substantially higher (median = 0.9, mean = 0.71; SD = 0.27), indicating strong similarity in their binding modes.

This similarity arises because, despite differences in their overall scaffolds, the ligands share common binding fragments that are positioned similarly within the active site. In particular, hydrogen bond acceptor groups (e.g., cyanophenyl and benzoxazinone moieties) are consistently oriented to form hydrogen bonds with the backbone of Ala270. The remainder of the binding pocket is dominated by an aromatic cage formed by Trp236, Tyr233, and Phe266 side chains [49], which further constrains ligand orientation and promotes similar interaction patterns.

### Binding sites and binding modes of ATP in circadian clock oscillator protein KaiC (Q79PF4)

Out of all the ATP interacting proteins within our data set, KaiC (UniProt ID: Q79PF4) had the largest number of ATP-binding complexes (256). It is one of the enzymatically active proteins in the KaiABC oscillator complex which controls the circadian clock in cyanobacteria [50]. KaiC is a homo-hexamer featuring KaiCI and KaiCII domains, and forms a double ring structure [50]. ATP molecules are wedged between subunits of KaiC hexamers with two distinct binding sites in CI and CII domains [50]. The binding modes of ATP in both domains also show distinctive differences. The structural studies of ATP have shown that the adenosine nucleobase only interacts with the protein in the CI domain, and no interaction occurs at the CII binding site, conferring high specificity for ATP over GTP in the CI domain [50].

Using the protein-ligand interaction data and clustering of ATPs’ binding site fingerprints, we were able to identify two distinct binding sites in the CI and CII domains (Figure 5 a). Similarly, clustering protein-ligand models by ECIFP similarities resulted in three clusters while using a cutoff distance of 0.38, determined based on the largest jump in linkage height observed in the hierarchical clustering dendrogram (Methods). Two of these clusters corresponded to the distinctive ATP binding modes in CI and CII domains (Figure 5 b). The third cluster had a single protein-ATP complex (PDB ID: 7v3x, auth_asym_id: X, auth_seq_id: 702). However, on visual inspection the binding mode of this ligand was similar to ATPs in CI domain, and would be clustered together with them if a slightly higher distance (0.4 or above) is used as cutoff. To better understand the differences between these ATP binding modes, we examined the enrichment of ligand interaction fingerprints (IFPs) using log odd ratio. The ligand IFP features correspond to the fragments of ligands near interacting atoms, hence enrichment of a set of fragments in one binding mode with respect to another indicates the likely difference in the pattern of protein-ligand model interactions. The top three ligand IFPs that are highly enriched in cluster 2 as compared to cluster 1 (log odd ratio < 0) included adenosine nucleobase, which was reported to interact strongly with CI domain compared to CII domain [50].

Distinctive ATP binding sites from each domain are coloured in cyan and blue sticks. **b,** Dendrogram of clusters formed based on ECIFP similarities of protein-ligand complexes - corresponding to two distinctive binding modes of ATP in CI and CII domains, coloured in cyan and blue. **c,** Enriched ECFP4 fragments of ATP interacting in the CI domain relative to CII domain are highlighted in red.

## Discussion

Analysis of protein-ligand interactions from experimentally resolved structures can provide valuable insights into the mechanisms underlying ligand recognition. The increasing availability of ligand-bound structures in the PDB offers an opportunity to systematically explore patterns of protein-ligand interactions. However, such analyses require scalable computational approaches. To address this, we propose ECIFP, an interaction-based fingerprint that enables comparison of ligand binding modes. Benchmarking results demonstrated that ECIFP outperforms ligand-only fingerprints in identifying similar binding sites with identical sequences but occupied by structurally diverse ligands. We further demonstrated its utility for identifying ligand binding modes across multiple PDB entries and comparing the binding modes of diverse ligands within the same binding site.

ECIFP builds on the concept of hashing atomic environments from the Extended Connectivity Fingerprint (ECFP) algorithm [51]. Unlike the original ECFP algorithm, which encodes all atoms of a molecule, ECIFP considers only the interacting atoms within a protein–ligand complex. Other interaction-based fingerprints based on ECFP, such as SPLIF [14] and PLEC [15] generate a single fingerprint representing the entire protein–ligand complex. While these approaches have been used for binding affinity prediction and docking pose scoring, collapsing ligand and protein interactions into a single fingerprint reduces interpretability when comparing interaction patterns across complexes. ECIFP addresses this limitation by generating separate fingerprints for the ligand and the protein, thereby enabling direct comparisons of ligand–ligand and protein–protein interactions between complexes. Furthermore, by analysing enriched fingerprint features and the atomic environments of contributing ligand atoms, ECIFP enables interpretation of differences in interaction patterns among different groups of complexes.

Despite its performance in detecting binding mode similarities, there remain areas for improvement. ECIFP currently generates binary fingerprints, ignoring interaction frequency and interaction types. This results in high ECIFP similarity for protein-ligand complexes dominated by non-specific hydrophobic interactions, but differ in their binding modes. The metric used for calculating the similarity of ECIFP is based on Tanimoto coefficients, and they are sensitive to ligand size [52]. For small ligands, even minor variations in interaction patterns can produce large differences in their ECIFP similarities. Future developments will aim to address these limitations. The ECIFPs of protein-ligand complexes and ligand interacting protein residues used to generate binding site fingerprints in this study are available from PDBe FTP (https://ftp.ebi.ac.uk/pub/databases/msd/pdbechem_v2/additional_data/interaction_fingerprints/). They can be used to calculate similarity of binding modes of ligands and group ligands binding to the same protein to distinct binding sites using the python code available from https://github.com/PDBeurope/ecifp. A key next step is to develop a graphical user interface (GUI) for exploiting this data and make it available via PDBe-KB Ligand pages [24] and PDBe-KB Protein pages [7].

## Methods

### Data set of protein-ligand complexes

For each PDB entry, its biological assemblies were generated from the asymmetric unit using the model-server [53], based on symmetry operations defined by the PDB entry authors or generated with PISA [54]. When multiple assemblies were available, we selected the smallest assembly containing all unique polymer chains (preferred assembly) [55]. For atoms with alternative locations, only the highest-occupancy conformer was retained.

Hydrogen atoms were added to ligands and proteins using ChimeraX [56]. Unique ligands were identified using PDBe CCDUtils [27], including cases where ligands consisted of multiple chemical components. protein–ligand interactions were then calculated using PDBe Arpeggio [23, 24].

### Quality filtering of protein-ligand complexes

For both binding mode and binding site analysis, ligands which are not likely to play biologically relevant roles were excluded [30, 31]. Ions, very small or large ligands with any of the following features were removed from binding mode analysis: number of heavy atoms less than twelve, molecular weight greater than 910 Da or the number of rings is zero. Ions were excluded from binding-site analysis when their total count in a protein exceeded 20, and carbohydrates were also removed as they were predominantly found to bind on the protein surface. Ligands were not filtered by quality metrics for binding site analysis, whereas for binding mode analysis, they were filtered using quality metrics defined in the wwPDB validation report [32, 33] (Figure 4 a.)

We included the ligand quality metrics and not for interacting residues for simplicity. For X-ray structures, PDB entries with resolution better than 4 Å were retained. Among these, PDB entries containing ligands with Real Space Correlation Coefficient (RSCC) > 0.9 were selected as high quality. PDB entries not meeting the 0.9 cutoff but with RSCC > 0.7 were further examined on their stereochemical quality. Similarly, PDB entries without reported RSCC values were also examined for stereochemical quality, if their resolution was better than 3 Å.

The stereochemical quality metrics in the wwPDB validation report include number of bond length outliers, number of bond angle outliers, bond RMSZ and angle RMSZ values. To find an acceptable number of bond length outliers and bond angle outliers for the ligands in the PDB, we selected a gold-standard protein ligand model by including only structures with resolution better than 2.5 Å and RSCC greater than 0.9. Using these gold-standard protein-ligand models, we defined the 95^th^ percentile as the cutoff value of bond length and bond angle outlier if the number of gold standard protein ligand models was less than 100. Otherwise, we performed random bootstrap sampling 1000 times and used the upper bound of 95^th^ percentile with 95% confidence interval as the outlier cutoff.

PDB entries with ligands with gold-standard protein-ligand models were then filtered using the bond angle and bond length outlier cutoff, and others were filtered based on the bond angle and bond length RMSZ (Root Mean Square Z score) cutoff of 2. For 3DEM PDB entries, a similar quality filtering procedure was followed, except we used a resolution of 7 Å and Q_Score based filtering as suggested by Pintilie et al. [57]. We did not include any NMR structures in our analysis, as they were only one percent of the total number of PDB entries with ligands.

### Ligand binding modes

To create fingerprint representations of protein-ligand models, the intermolecular interactions between protein-ligand pairs were calculated using PDBe Arpeggio [23, 24]. PDBe Arpeggio calculates atom-atom, atom-plane and plane-plane interactions based on atom types and distance cutoffs [23]. Out of the 15 types of atom-atom interactions calculated by PDBe Arpeggio, we removed some of the non-specific interactions including proximal, clash, VdW (Van der Waals) clash and VdW interactions. For the plane-plane interactions, all the atoms which are part of the planes are considered as interacting atoms. PDBe CCDUtils [27] was used to parse the ligand and amino acid residues’ coordinate data in CIF files [25] and generated ECFP4 fingerprints of length 1024 using RDKit [58].

Given two protein-ligand complexes A and B with ligand IFPs *l_A_*and *l_B_*, and protein IFPs *p_A_* and *p_B_*, the similarity between them were calculated as the weighted geometric mean of Tanimoto coefficient [59] between ligand IFPs and protein IFPs as in equation 1. For binding mode comparison of different ligands within the same binding site, we used *w_l_* = 0.6 and *w_p_* = 0.4, giving higher weight for ligand IFP similarity. Conversely, for comparing the same ligand across different binding sites, *w*□ = 0.4 and *w*□ = 0.6 were applied to prioritise protein IFP similarity. In both cases, weights were constrained to sum to one to ensure a balanced similarity measure while reflecting the expected relative contributions of ligand and protein interactions.

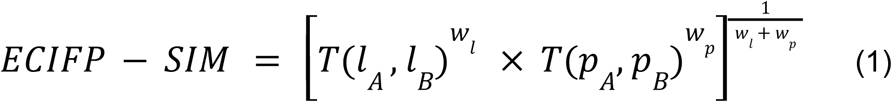

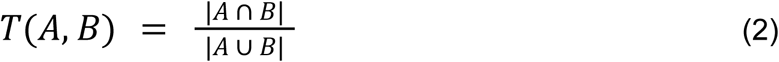

The ECIFP similarity values range between 0 and 1, with 1 being identical binding modes. After calculating the pairwise similarities between the protein-ligand complexes, the similarity values were converted to distance (1-similarity). Pairwise distance values were used to perform hierarchical clustering using the minimax linkage method [34]. We selected the optimal distance cutoff based on the largest jump in linkage height observed in the hierarchical clustering dendrogram, because a large jump typically indicates that significantly different clusters are getting merged. Specifically, we identified the point at which the largest increase in linkage height occurs, provided that the corresponding distance is greater than 0.25 and less than or equal to 0.6. If no such point was found, a default cutoff of 0.5 was used. A maximum distance cutoff of 0.6 was used to avoid clusters containing highly dissimilar data points. Similarly, a minimum distance cutoff was chosen to avoid sparse clusters. We used PyMinimax [60] for minmax linkage matrix computation and SciPy [61] for clustering. To identify enriched fragments of ligands between a pair of clusters, Fisher’s exact test was done on the contingency table of each ligand IFP feature using SciPy [61] and then corrected p-values for multiple testing using Benjamini/Hochberg method [62] using statsmodel [63]. Using a significance level of 0.05 for the corrected p-values, we selected statistically significant ligand IFPs and identified the enriched IFP using the log odd ratio.

### Binding site similarity assessment

The benchmark data used for binding site similarity assessment was a subset of ProSPECCTs [22] data set available from http://www.ewit.ccb.tu-dortmund.de/ag-koch/prospeccts/. We used the *kahraman_structures<u>.tar.gz</u>, identical_structures<u>.tar.gz</u>*, *barelier_structures<u>.tar.gz</u>*, and *review_structures<u>.tar.gz</u>* for this study. Each of these folders contained a CSV file with pairs of protein-ligand complexes with their classes *active* for similar and *inactive* for dissimilar binding sites, and a compressed folder with PDB files of protein-ligand complexes. The mapping of ligands between our data set and ProSPECCTs was done based on a combination of *entry_id, auth_asym_id* and *auth_seq_id* fields in the mmCIF files. As protein-ligand complexes were represented using only *entry_id* and *auth_asym_id* in the CSV files available from ProSPECCTs, we extracted the *auth_seq_id* information from the supplementary tables of ProSPECCTs paper for *barelier_structures* and *review_structures.* For other data sets, we extracted the *auth_seq_id* from the PDB files by parsing them using Gemmi [64].

For ECIFP similarity, we used an equal weight of 0.5 for both ligand IFP and protein IFP. A cutoff value of ECIFP similarity for the classification task was determined by fitting a gamma distribution function to the non-zero ECIFP similarity values of randomly selected 10,000,000 protein-ligand model pairs and selecting the 95th percentile.

### Ligand binding sites

For each protein, the distinctive binding sites were identified by clustering protein-ligand complexes based on the similarity of binding fingerprints using Szymkiewicz–Simpson coefficient [65]. The binding fingerprints were defined as the set of UniProt residue numbers corresponding to ligand-contacting residues [47]. We included proteins with more than one instance of interacting ligand with at least 3 interacting residues through their side chain atoms in our analysis. Additionally, in case of proteins with proteolytic cleavage products, each of them were clustered separately. For clustering, we used the agglomerative hierarchical clustering algorithm with the single linkage method implementation in SciPy [54], and distance cutoff of 0.4 was used for all the proteins. For each ligand cluster, the set of all amino acids interacting with at least 25% of the ligands in the cluster were considered as the binding site residues. To evaluate the quality of binding site clustering, the minimum interatomic distance between ligands occupying different binding sites of the same protein was calculated following structural superposition. For each protein, the preferred assemblies [55] of all corresponding structures were aligned to a representative structure using USalign [40] with the oligomer alignment option enabled. The representative structure was selected as the PDB entry with highest sequence coverage along with better resolution [7].

## Data and Software Availability

The data used and generated in the analysis are available from Zenodo (https://doi.org/10.5281/zenodo.18557398). The ECIFPs of protein-ligand complexes and ligand interacting protein residues are available from PDBe FTP (https://ftp.ebi.ac.uk/pub/databases/msd/pdbechem_v2/additional_data/interaction_fingerprints/). The Python code for the developed method and for reproducing the results reported in this study is publicly available on GitHub: https://github.com/PDBeurope/ecifp

## Supporting Information

Additional results including validation of ECIFP for binding site comparison (PDF)

## Author Contributions

Conceptualization: IRK, PC and SV; methodology: IRK; Python code for data processing and method implementation: IRK; writing—original draft preparation: IRK; writing—review and editing: PC, JF, SV; visualization: AM; Project Administration: PC, infrastructure and data access: BB, SS; supervision: PC, SV; funding acquisition: SV. All authors have read and agreed to the published version of the manuscript.

