## Supporting Information for "Large-scale analysis of ligand binding mode similarities in the PDB using interaction fingerprints"

| CCD ID | Name | Number of complexes | Number of binding modes |
|---|---|---|---|
| FAD | Flavin adenine dinucleotide | 4889 | 39 |
| NAD | Nicotinamide-adenine dinucleotide | 4706 | 37 |
| FMN | Flavin Mononucleotide | 2621 | 17 |
| PLP | Pyridoxal 5'-phosphate | 2324 | 18 |
| COA | Coenzyme A | 1438 | 38 |
| H4B | Biopterin | 1056 | 3 |
| SAM | S-adenosylmethionine | 968 | 28 |
| GSH | Glutathione | 774 | 22 |
| BTN | Biotin | 282 | 12 |
| HEA | Heme | 240 | 2 |
| MGD | Molybdopterin | 182 | 2 |
| PNS | Phosphopantetheine | 79 | 6 |
| ASC | Ascorbic acid | 65 | 32 |
| PQQ | Pyrroloquinoline quinone | 58 | 2 |
| F43 | Factor F430 | 45 | 1 |

| | | | |
|---|---|---|---|
| TP7 | Coenzyme B | 45 | 1 |
| FFO | Tetrahydrofolic acid | 38 | 8 |
| MQ7 | Menaquinone | 127 | 2 |
| UQ1 | Ubiquinone | 19 | 11 |
| DPM | Dipyrromethane | 14 | 1 |
| LPA | Lipoic acid | 4 | 1 |

Supplementary Table 1. Binding modes of 21 organic cofactors defined in the CoFactor database [1], identified by clustering of ECIFP similarities. CCD ID corresponds to the ID of the cofactor in Chemical Component Dictionary (CCD) [2]. Number of complexes is the number of unique instances of cofactors in the subset of PDB used in this study. Number binding modes correspond to the number of clusters of each cofactor.

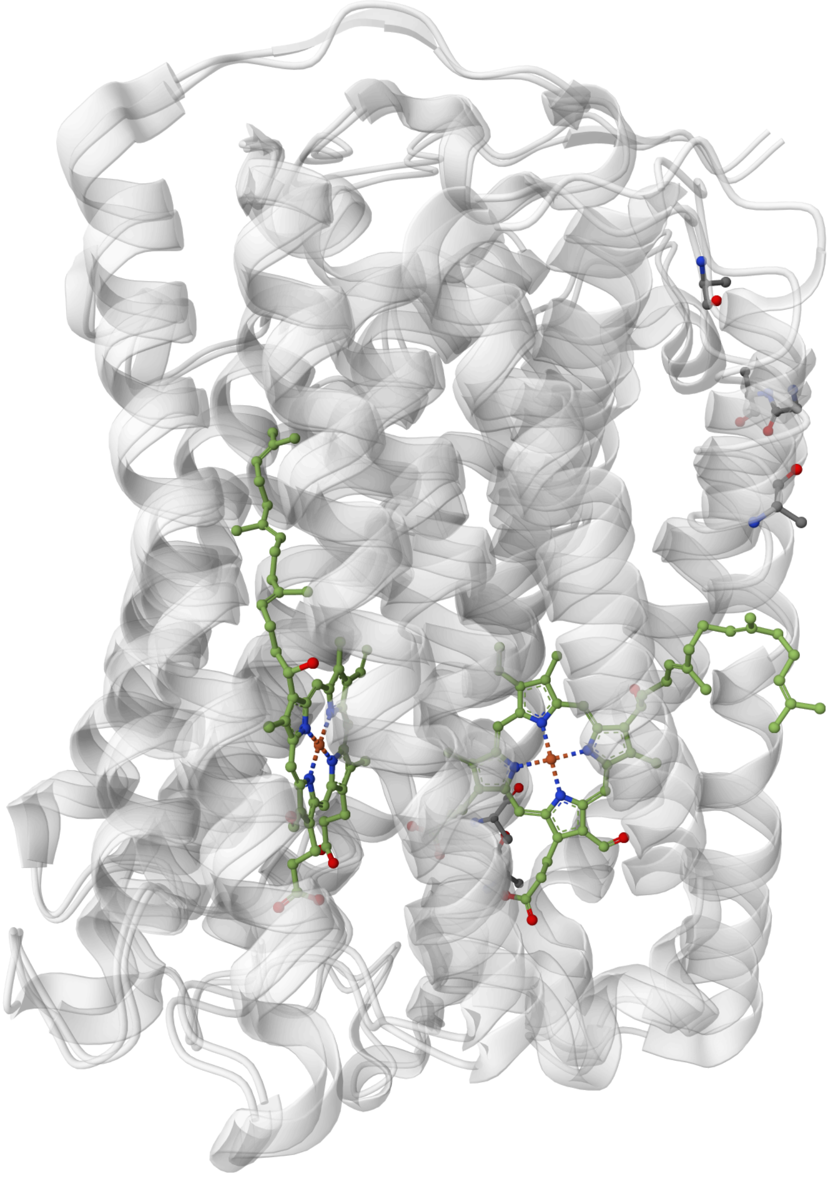

Supplementary Figure 1. Two different binding modes of HEME-A (HEA) in cytochrome c oxidase. Chain A and chain BN from PDB entries 6nmf and 5j4z are aligned using US-align [3] and visualised using Mol* viewer [4]. ECIFP similarities between these protein-HEA complexes were 0.323

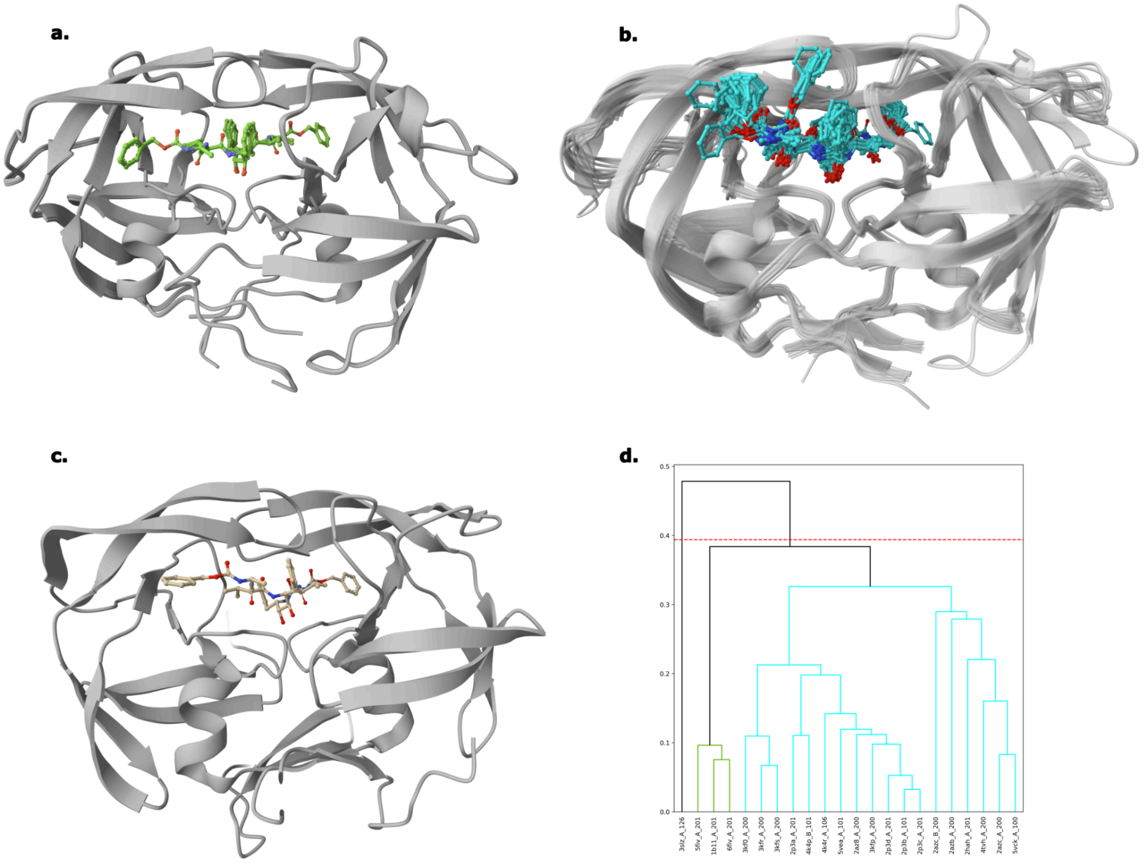


Supplementary Figure 2. Three distinct binding modes of retroviral protease inhibitor TL-3 identified by clustering. **a**, Superposed view of cluster one with TL3 bound to structures of feline immunodeficiency virus proteases shown in green sticks. **b**, Superposed view of cluster two with TL3 bound to structures of human immunodeficiency virus proteases in cyan sticks. **c**, Cluster three with TL3 bound to the structure of murine leukemia virus-related virus protease in wheat tint stick. **d**, Dendrogram showing the hierarchical clusters, coloured according to cluster membership

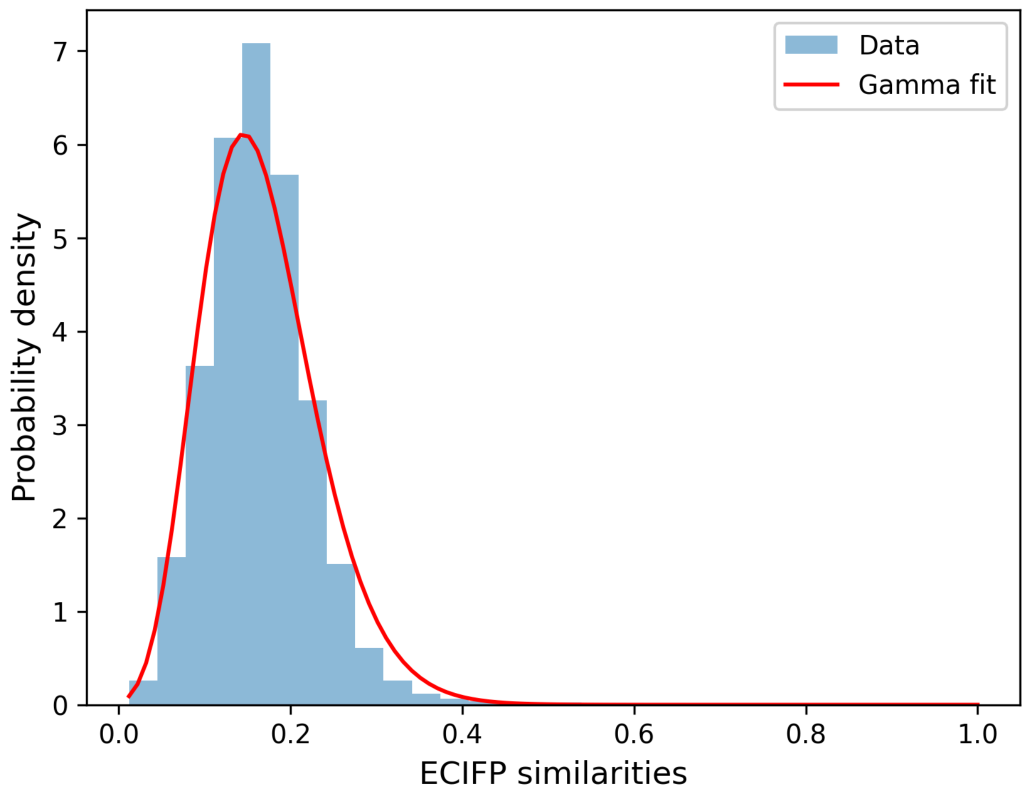


Supplementary Figure 3. Distribution of ECIFP similarity scores for 10,000,000 randomly selected protein–ligand complex pairs. The fitted Gamma distribution is shown in red. The 95th percentile of the fitted Gamma distribution of ECIFP similarities corresponds to a value of 0.289.

| | Precision | | Recall | | F1 score | |
|---|---|---|---|---|---|---|
| | Baseline | ECIFP | Baseline | ECIFP | Baseline | ECIFP |
| Structures with identical sequences (DS1) | 0.76 | 0.73 | 0.24 | 0.61 | 0.36 | 0.67 |
| Kahraman data set (DS2) | 0.31 | 0.35 | 1 | 0.98 | 0.48 | 0.52 |
| Barelier data set (DS3) | 0.20 | 0.22 | 0.89 | 0.88 | 0.33 | 0.35 |
| Data set of successful applications (DS4) | 0.001 | 0.001 | 0.41 | 0.41 | 0.003 | 0.003 |

Supplementary Table 2. Performance of ECIFP similarities for binding site comparison

| Pair 1 | | Pair 2 | | | | | |
|---|---|---|---|---|---|---|---|
| Identifier | Ligand | identifier | Ligand | Ligand IFP similarity | Protein IFP similarity | ECIFP similarity | Ligand ECFP4 similarity |
| 1bn1_A_555 | AL5 | 1bnv_A_555 | AL7 | 0.295 | 0.932 | 0.525 | 0.265 |
| 1bnn_A_555 | AL1 | 1bn1_A_555 | AL5 | 0.255 | 1 | 0.505 | 0.26 |
| 1bnn_A_555 | AL1 | 1bnw_A_555 | TPD | 0.295 | 1 | 0.544 | 0.253 |
| 1bnv_A_555 | AL7 | 1bnw_A_555 | TPD | 0.279 | 0.932 | 0.51 | 0.244 |
| 1cim_A_264 | PTS | 1i91_A_555 | INQ | 0.349 | 0.838 | 0.541 | 0.289 |
| 1i8z_A_555 | INL | 4bf6_A_1263 | X0Q | 0.302 | 0.87 | 0.512 | 0.25 |
| 1okl_A_862 | MNS | 1bnv_A_555 | AL7 | 0.268 | 0.953 | 0.506 | 0.231 |
| 1qiq_A_332 | ACC | 1odn_A_1324 | APV | 0.407 | 0.806 | 0.573 | 0.241 |
| 1qjf_A_1333 | ACS | 1qiq_A_332 | ACC | 0.481 | 0.821 | 0.629 | 0.291 |
| 1ype_H_5555 | UIP | 3rm2_H_1 | S00 | 0.325 | 0.897 | 0.54 | 0.294 |
| 1ypk_H_5555 | CCR | 1ype_H_5555 | UIP | 0.338 | 0.923 | 0.558 | 0.282 |
| 1ypk_H_5555 | CCR | 2zda_H_1000 | 32U | 0.306 | 0.829 | 0.503 | 0.288 |
| 2aw1_A_264 | COX | 2hl4_A_264 | BO1 | 0.333 | 0.784 | 0.511 | 0.263 |
| 2byh_A_1226 | 2D7 | 3bmy_A_1 | CXZ | 0.344 | 0.759 | 0.511 | 0.293 |
| 2byi_A_1226 | 2DD | 3bmy_A_1 | CXZ | 0.323 | 0.868 | 0.529 | 0.284 |
| 2hu6_A_400 | 37A | 3ts4_A_306 | EEG | 0.35 | 0.812 | 0.533 | 0.278 |
| 2hu6_A_400 | 37A | 3tsk_A_306 | QEG | 0.357 | 0.765 | 0.523 | 0.267 |
| 2hu6_A_400 | 37A | 4efs_A_306 | E37 | 0.295 | 0.862 | 0.504 | 0.28 |
| 3chp_A_901 | 4BO | 3fh5_A_611 | 24P | 0.317 | 0.904 | 0.535 | 0.297 |
| 3f17_A_0 | HS4 | 3tsk_A_306 | QEG | 0.34 | 0.754 | 0.506 | 0.291 |
| 3m04_A_502 | BE9 | 3m2x_A_502 | BEX | 0.34 | 0.792 | 0.519 | 0.267 |
| 3mhl_A_265 | J71 | 1kwq_A_900 | SG1 | 0.361 | 0.7 | 0.503 | 0.295 |
| 3mhl_A_265 | J71 | 2hl4_A_264 | BO1 | 0.351 | 0.804 | 0.532 | 0.274 |
| 3mhl_A_265 | J71 | 3mzc_A_263 | S6I | 0.371 | 0.721 | 0.517 | 0.269 |
| 3oys_A_263 | OYS | 4knj_A_302 | E1F | 0.275 | 0.918 | 0.502 | 0.296 |
| 4bf6_A_1263 | X0Q | 1bnm_A_555 | AL8 | 0.321 | 0.848 | 0.522 | 0.291 |
| 4bf6_A_1263 | X0Q | 1bnv_A_555 | AL7 | 0.327 | 0.932 | 0.552 | 0.291 |
| 4bf6_A_1263 | X0Q | 1bnw_A_555 | TPD | 0.292 | 0.872 | 0.504 | 0.29 |

Supplementary Table 3. Binding site pairs with class *active* in DS1, but having low ligand similarities and high ECIFP similarities.

| Pair 1 | | Pair 2 | | | | | |
|---|---|---|---|---|---|---|---|
| Identifier | Ligand | Identifier | Ligand | Ligand IFP similarity | Protein IFP similarity | ECIFP similarity | Ligand ECFP4 similarity |
| 12as_A_332 | AMP | 1ayl_A_544 | ATP | 0.69 | 0.585 | 0.636 | 0.831 |
| 12as_A_332 | AMP | 1b8a_A_500 | ATP | 0.763 | 0.56 | 0.654 | 0.831 |
| 12as_A_332 | AMP | 1e2q_A_302 | ATP | 0.73 | 0.442 | 0.568 | 0.831 |
| 12as_A_332 | AMP | 1esq_A_300 | ATP | 0.711 | 0.489 | 0.59 | 0.831 |
| 12as_A_332 | AMP | 1gn8_A_700 | ATP | 0.784 | 0.426 | 0.578 | 0.831 |
| 12as_A_332 | AMP | 1hex_A_400 | NAD | 0.75 | 0.35 | 0.512 | 0.584 |
| 1a0i_A_1 | ATP | 1amu_A_567 | AMP | 0.69 | 0.483 | 0.578 | 0.831 |
| 1a49_A_535 | ATP | 1kht_B_2193 | AMP | 0.667 | 0.385 | 0.506 | 0.831 |
| 1ayl_A_544 | ATP | 1jq5_A_401 | NAD | 0.534 | 0.492 | 0.513 | 0.644 |
| 1cq1_A_455 | BGC | 1k1w_A_660 | GLC | 0.368 | 0.758 | 0.528 | 1 |
| 1e8x_A_3000 | ATP | 1amu_A_567 | AMP | 0.659 | 0.417 | 0.524 | 0.831 |
| 1ej2_A_1339 | NAD | 1b8a_A_500 | ATP | 0.527 | 0.481 | 0.503 | 0.644 |
| 1gn8_A_700 | ATP | 1kht_B_2193 | AMP | 0.833 | 0.471 | 0.627 | 0.831 |
| 1hex_A_400 | NAD | 1b8a_A_500 | ATP | 0.659 | 0.459 | 0.55 | 0.644 |
| 1hex_A_400 | NAD | 1tid_A_200 | ATP | 0.558 | 0.494 | 0.525 | 0.644 |
| 1kht_B_2193 | AMP | 1b8a_A_500 | ATP | 0.718 | 0.446 | 0.566 | 0.831 |
| 1kvk_A_535 | ATP | 1amu_A_567 | AMP | 0.705 | 0.519 | 0.604 | 0.831 |
| 1mew_A_987 | NAD | 1b8a_A_500 | ATP | 0.58 | 0.433 | 0.501 | 0.644 |
| 1qb8_A_300 | AMP | 1tid_A_200 | ATP | 0.8 | 0.527 | 0.649 | 0.831 |
| 1rlz_A_700 | NAD | 1b8a_A_500 | ATP | 0.63 | 0.583 | 0.606 | 0.644 |
| 1rlz_A_700 | NAD | 1gn8_A_700 | ATP | 0.526 | 0.486 | 0.506 | 0.644 |
| 1rlz_A_700 | NAD | 1tid_A_200 | ATP | 0.627 | 0.571 | 0.599 | 0.644 |
| 1s7g_B_701 | NAD | 1tid_A_200 | ATP | 0.526 | 0.494 | 0.51 | 0.644 |
| 1tid_A_200 | ATP | 1jp4_A_601 | AMP | 0.6 | 0.539 | 0.569 | 0.831 |
| 1tid_A_200 | ATP | 1kht_B_2193 | AMP | 0.705 | 0.538 | 0.616 | 0.831 |
| 2a5f_B_1536 | NAD | 1ayl_A_544 | ATP | 0.545 | 0.459 | 0.5 | 0.644 |
| 2gbp_A_310 | BGC | 1k1w_A_660 | GLC | 0.857 | 0.385 | 0.574 | 1 |
| 2npx_A_818 | NAD | 1amu_A_567 | AMP | 0.569 | 0.5 | 0.533 | 0.584 |

Supplementary Table 4. Binding site pairs with class *inactive* in DS2, but having high ECIFP similarities

| Pair 1 | | Pair 2 | | | | | |
|---|---|---|---|---|---|---|---|
| Identifier | Ligand | identifier | Ligand | Ligand IFP similarity | Protein IFP similarity | ECIFP similarity | Ligand ECFP4 similarity |
| 1e6w_A_301 | NAD | 1e3w_D_301 | NAD | 0.877 | 0.962 | 0.919 | 1 |
| 1gvr_A_401 | FMN | 3p74_A_401 | FMN | 0.9 | 0.79 | 0.843 | 1 |
| 1gal_A_600 | FAD | 1gpe_B_600 | FAD | 0.845 | 0.656 | 0.744 | 1 |
| 1xel_A_340 | NAD | 1orr_B_1300 | NAD | 0.95 | 0.578 | 0.741 | 1 |
| 1xel_A_340 | NAD | 2yy7_A_3001 | NAD | 0.919 | 0.597 | 0.741 | 1 |
| 2hs6_A_401 | FMN | 1vhn_A_322 | FMN | 0.968 | 0.537 | 0.721 | 1 |
| 1xel_A_340 | NAD | 1e3w_D_301 | NAD | 0.935 | 0.534 | 0.707 | 1 |
| 2hs6_A_401 | FMN | 3p74_A_401 | FMN | 0.794 | 0.606 | 0.694 | 1 |
| 1xel_A_340 | NAD | 2b69_A_800 | NAD | 0.902 | 0.507 | 0.676 | 1 |
| 1e6w_A_301 | NAD | 3orf_A_901 | NAD | 0.774 | 0.571 | 0.665 | 1 |
| 1e6w_A_301 | NAD | 2b69_A_800 | NAD | 0.844 | 0.524 | 0.665 | 1 |
| 1z41_A_1500 | FMN | 3l5l_A_1401 | FMN | 0.844 | 0.506 | 0.654 | 1 |
| 1xel_A_340 | NAD | 3orf_A_901 | NAD | 0.831 | 0.513 | 0.653 | 1 |
| 2hs6_A_401 | FMN | 3l5l_A_1401 | FMN | 0.871 | 0.489 | 0.652 | 1 |
| 1xel_A_340 | NAD | 2gn8_B_334 | NAP | 0.836 | 0.506 | 0.651 | 0.88 |
| 1gal_A_600 | FAD | 3ad9_B_405 | FAD | 0.855 | 0.49 | 0.647 | 1 |
| 1xel_A_340 | NAD | 1u1i_C_1196 | NAD | 0.919 | 0.449 | 0.642 | 1 |
| 1gal_A_600 | FAD | 3vqr_B_1001 | FAD | 0.853 | 0.479 | 0.639 | 1 |
| 1e6w_A_301 | NAD | 2yy7_A_3001 | NAD | 0.862 | 0.474 | 0.639 | 1 |
| 1e6w_A_301 | NAD | 1xg5_D_1304 | NAP | 0.791 | 0.514 | 0.638 | 0.88 |
| 1e6w_A_301 | NAD | 1u1i_C_1196 | NAD | 0.862 | 0.458 | 0.628 | 1 |
| 1gal_A_600 | FAD | 4at0_A_1493 | FAD | 0.951 | 0.413 | 0.627 | 1 |
| 1gvr_A_401 | FMN | 1vhn_A_322 | FMN | 0.9 | 0.43 | 0.622 | 1 |
| 1xel_A_340 | NAD | 3cin_A_400 | NAD | 0.869 | 0.446 | 0.622 | 1 |
| 1gal_A_600 | FAD | 3g5s_A_444 | FAD | 0.732 | 0.52 | 0.617 | 1 |
| 1xel_A_340 | NAD | 2yyy_A_2001 | NAP | 0.762 | 0.5 | 0.617 | 0.88 |
| 1xel_A_340 | NAD | 1xg5_D_1304 | NAP | 0.815 | 0.462 | 0.614 | 0.88 |
| 1gvr_A_401 | FMN | 3l5l_A_1401 | FMN | 1 | 0.376 | 0.613 | 1 |
| 1gal_A_600 | FAD | 1c0k_A_1363 | FAD | 0.746 | 0.5 | 0.611 | 1 |
| 1e6w_A_301 | NAD | 4idg_A_402 | NAD | 0.855 | 0.435 | 0.61 | 1 |
| 1gal_A_600 | FAD | 4b68_A_1492 | FAD | 0.783 | 0.471 | 0.607 | 1 |

| | | | | | | | |
|---|---|---|---|---|---|---|---|
| 1gal_A_600 | FAD | 3ka7_A_500 | FAD | 0.841 | 0.438 | 0.606 | 1 |
| 1xel_A_340 | NAD | 4idg_A_402 | NAD | 0.823 | 0.446 | 0.606 | 1 |
| 1gal_A_600 | FAD | 2yr5_B_801 | FAD | 0.868 | 0.417 | 0.602 | 1 |

Supplementary Table 5. Binding site pairs with class *inactive* in DS4, but having high ECIFP similarities

| Protein name | Uniprot ID | Number of binding sites | Remarks |
|---|---|---|---|
| Prostaglandin G/H synthase 2 | Q05769 | 2 | Clusters the substrate binding site and cofactor binding site separately |
| Aminoglycoside phosphotransferase | Q47396 | 2 | Clusters the substrate binding site and cofactor binding site separately |
| Phenylethanolamine N-methyltransferase | P11086 | 1 | Groups all the ligands in the substrate binding site to the same cluster |
| Nuclear receptor ROR-gamma | P51449 | 4 | Each of the binding sites are distinct in the 3D view |
| Muscarinic acetylcholine receptor M2 | P08172 | 2 | Clustered the orthosteric and allosteric ligand binding sites separately |
| Tyrosine-protein kinase ABL1 | P00519 | 5 | Identifies four distinct ligand binding clusters. One binding site is occupied by ion NA |
| Glucose-6-phosphate 1-dehydrogenase | P11413 | 3 | Three distinct ligand binding sites were found in 3D superposed view |
| Kinesin-like protein KIF11 | P52732 | 3 | Three distinct ligand binding sites were found in 3D superposed view |
| Dihydrofolate reductase | C3TR70 | 2 | Two distinct ligand binding sites were found in 3D superposed view |
| Clostridium Botulinum C3 Exoenzyme | P15879 | 1 | One binding site occupied by ligands was found in the 3D superposed view |
| L-arabinose-binding protein | P02924 | 1 | One binding site occupied by ligands was found in the 3D superposed view |
| Dihydropteroate synthase | P0AC13 | 1 | One binding site occupied by ligands was found in the 3D superposed view |
| Glucokinase | P35557 | 3 | Three distinct ligand binding sites were found in 3D superposed view |
| RNMT | O43148 | 1 | One binding site occupied by ligands was found in the 3D superposed view |
| Fatty acid binding protein | Q01469 | 2 | Two distinct ligand binding sites were found in 3D superposed view |
| Vitamin D3 receptor A | Q9PTN2 | 2 | Two distinct ligand binding sites were found in 3D superposed view |

| | | | |
|---|---|---|---|
| Beta-galactoside alpha-2,6-sialyltransferase 1 | P15907 | 2 | Can be considered as a single binding site. One of ligand occupies both sites |
| Beta-1,4-galactosyltransferase 1 | P15291 | 2 | Two distinct ligand binding sites were found in 3D superposed view |
| MoeE5 | A0A003 | 2 | Two distinct ligand binding sites were found in 3D superposed view |
| Adenylate cyclase | X8CHM4 | 1 | One binding site occupied by ligands was found in the 3D superposed view |

Supplementary Table 6. Number of binding sites identified for 20 randomly selected proteins
